# Oscillatory correlates of visuomotor control under varying amount of feedback delay

**DOI:** 10.64898/2026.06.04.729854

**Authors:** Peng Wang, Josephine Gräfe, Zhenyu Wang, Fanni Peters, Jonathan Yi, Jakub Limanowski

## Abstract

A weighted integration of visual and proprioceptive movement feedback is key for an adaptive body representation by the brain. Previous work has suggested a relationship between alpha and beta oscillations in sensory cortices and inter-sensory attentional control under visuo-proprioceptive incongruence compared with congruence; and frontal theta oscillations as a marker of the related cognitive-attentional control and response inhibition. Here, we asked whether these oscillatory correlates indicate a gradual adjustment of attention and the corresponding sensory weights, rather than merely a response to conflict per se. Thus, we adopted a recently developed virtual reality based hand-target phase matching task: Participants had to track a target oscillation with recurrent grasping movements of a virtual hand, which they controlled via a data glove. We added a small or a large delay to the virtual hand’s movements, introducing varying amounts of visuo-proprioceptive conflict. Concurrent EEG recordings revealed a delay-dependent modulation of (i) sensorimotor beta oscillations, (ii) oscillations “entrained” at key movement frequencies over central-parietal sensors, and (iii) mid-frontal theta oscillations. Granger causality analysis suggested that entrained oscillations were more strongly predictive of visual kinematic signals, whereas the prediction of somatosensory kinematics was attenuated depending on the amount of delay. Finally, phase-amplitude couplings suggested that sensorimotor beta coupled with the movement-related oscillations as well as mid-frontal theta. Our results thus establish key oscillatory correlates that scale with the amount of visuo-proprioceptive conflict during visually guided action.

## 1. Introduction

A weighted integration of visual and proprioceptive data is key for guiding movements of the upper extremities (Sober & Sabes, 2005; van Beers et al., 1999; Ernst & Banks, 2002; Graziano, 1999; Avillac et al., 2007). The underlying processes can be studied by artificially introducing conflicts between the two modalities, for instance through mirror-reversal or by adding a temporal lag to the visual movement feedback. This line of work suggests that, when vision is task relevant but conflicting with proprioception, visual movement signals are augmented, while proprioceptive signals may be temporarily attenuated (Bernier et al., 2009; Balslev et al., 2004; Zeller et al., 2015; see Moorselaar & Slagter, 2020; Limanowski, 2022, for reviews). In principle, this can be interpreted as the result of inter-sensory re-weighting driven “top-down” by cognitive-attentional factors (Kelso et al., 1975; Rohe & Noppeney, 2018; González-García et al., 2020; Limanowski & Friston, 2020).

The neural correlates of these processes may be illuminated through the study of cortical oscillations as measured with M/EEG, as an index of large-scale neuronal communication (Schroeder & Lakatos, 2009; Clayton et al., 2015; Senkowski & Engel, 2024; Kaltenmaier et al., 2025). It is generally assumed that rhythmic fluctuations in neuronal excitability temporally structure information transfer within and between cortical regions. In hierarchical models of cortical communication, distinct frequency bands are thought to support different directions of information flow, thereby enabling flexible routing of “bottom-up” (sensory) and “top-down” (e.g., cognitive-attentional) signals (Fries 2015; Bastos et al., 2015; Michalareas et al., 2016; Miller; Spitzer & Haegens, 2017; Arnal et al., 2011; Senkowski & Engel, 2024).

In particular, several low-frequency oscillatory bands have been identified as potential markers of cognitive-attentional control over sensory processing and multisensory integration: Directing endogenous attention to a specific sensory modality has been linked to the attenuation of corresponding sensory low-frequency oscillatory power in the alpha and (low) beta ranges in visual (Thut et al., 2006; Foxe & Snyder, 2011; Jensen & Mazaheri, 2010), auditory (Foxe et al., 1998; Weisz et al., 2011; Gomez-Ramirez et al., 2011), and somatosensory tasks (Bauer et al., 2012a,b; van Ede et al., 2010, 2011; Anderson & Ding, 2011; Haegens et al., 2011; Zeller et al., 2015; Jones et al., 2010). Beta oscillations have furthermore been linked to the integration of (conflicting) multisensory information (Senkowski & Engel, 2024; Arnal et al., 2011; Keil et al., 2016) and sensorimotor coordination (Morillon & Baillet, 2017; Saleh et al., 2010; Arnal et al., 2014; Fujioka et al., 2015; Kulashekhar et al., 2016; Pollok et al., 2008). Interestingly, these oscillations may be cyclically modulated: Thus, the amplitude of alpha/beta oscillations can be “entrained” i.e. modulated by rhythmic stimuli or body movements, potentially suggesting an optimized allocation of resources to behaviorally relevant time points (Gomez-Ramirez et al., 2011; Wang & Limanowski, 2023). Another key marker of endogenous, selective attention and cognitive control in particular during inter-sensory conflicts are theta oscillations recorded over mid-frontal regions (Ullsperger et al., 2014; Cavanagh and Frank 2014; Cohen and Donner 2013; Zavala et al., 2018; Murray et al., 2025; Spooner et al., 2019). A coupling of theta and alpha/beta oscillations has likewise been suggested as a potential mechanism for top-down control (Chacko et al., 2018; Watanabe et al., 2021; Wendiggensen et al., 2023).

In sum, there is good reason to assume that low-frequency oscillations indicate the effects of “top-down” context-dependent weighting and integration of (conflicting) visual and proprioceptive movement information during action (cf. Balslev et al., 2004; Bernier et al., 2009; Press et al., 2011; Lebar et al., 2017). However, previous designs comparing incongruent with congruent visual movement feedback cannot with certainty answer whether these oscillatory correlates indeed reflected a proportional adjustment of e.g. sensory weights and cognitive-attentional control, or perhaps some more general, qualitative difference related to moving under synchronous vs asynchronous visual and proprioceptive feedback.

Therefore, in the present EEG study, we tested for oscillatory differences related to moving under varying degrees of visuo-proprioceptive conflict. We adopted a previously established hand-target phase matching task, where participants have to align the grasping rhythm of their real, unseen hand or a visible, delayed virtual hand with a target rhythm (Limanowski & Friston, 2020; Limanowski et al., 2020). Note that this task was designed as a sustained, continuous-movement task under inter-sensory conflict (cf. Kelso et al., 1975; Rohe & Noppeney, 2018; González-García et al., 2020), not as a visuomotor adaptation task. I.e., as participants were trained extensively on the visual feedback delays, we did not expect or target sensorimotor learning. Instead, we focused on the cognitive-attentional processes related to using visual movement feedback that conflicted with proprioception for continuous (periodic) goal-directed action. Importantly, here, instead of comparing tracking under delayed vs synchronous visual movement feedback (i.e., implying visuo-proprioceptive conflict vs no-conflict), we introduced two levels of visuomotor delay (small vs large, which determined the amount of visuo-proprioceptive conflict). Furthermore, we orthogonally manipulated the target frequency (0.3 vs 0.5 Hz, which sets the expected timing of grasp transitions) to replicate delay dependent effects across different movement frequencies. We thereby assumed that the underlying control and re-weighting processes—indicated by low-frequency (alpha/beta-band) dynamics—should be frequency-invariant (present for both slow and fast grasping), whereas the magnitude of these signals should scale with visuo-proprioceptive conflict (i.e., showing a delay main effect).

We first tested for conflict (i.e., delay) dependent spectral effects at the key target frequency and its first harmonic (potentially related to “entrainment”; cf. Gomez-Ramirez et al., 2011; Saleh et al., 2010; Arnal et al., 2025; Wang & Limanowski, 2023). Then, we applied spectral Granger prediction to test for a potential predictive coupling of EEG signals and (hand) movement signals at electrodes showing delay-dependent modulation at these frequencies. Thus, we aimed to clarify whether the modulation of oscillatory EEG power at movement frequencies could be interpreted as predictive of hand kinematics, or vice versa. Finally, our spectral analysis revealed delay effects on the power of beta oscillations (likely in somatosensory regions), and on parietal oscillations at their modulating movement frequencies, and midfrontal theta oscillations. As the latter two could, potentially, control signals from posterior parietal or prefrontal regions (Senkowski et al., 2008; Buschmann & Miller, 2007; Ullsperger et al., 2014), we also tested for phase-amplitude coupling between theta-beta and f1-beta both within and across the identified sites.

## 2. Methods

### 2.1 Experimental setup

#### 2.1.1 Participants

26 participants (20 females, aged 19-31, mean±SD = 22±3 years), who successfully completed a prior training session, participated in the experiment. The target sample size of 26 participants was determined using a power analysis based on effect sizes from our previous study (Wang & Limanowski, 2023), which tested for phase–amplitude coupling (PAC) between grasping phase and beta power using a Wilcoxon signed-rank test. In that study (18 participants), the effect size was r = 0.6868 (signed-rank statistic = 153, z = 2.92, ɑ = 0.05), corresponding to a statistical power of 0.86. Increasing the desired power to 0.95 yielded a required sample size of 26 participants (calculated using G*Power (Faul et al., 2007)). All participants were right-handed, had normal or corrected-to-normal vision, and reported no history of psychiatric or neurological conditions. They received 10€/hour or student credit as compensation. Written informed consent was obtained from all participants prior to participation.

#### 2.1.2 Statement of ethical approval

This study was approved by the local Ethics Committee of the University Medicine of Greifswald and performed in accordance with this approval, and the relevant guidelines and regulations.

#### 2.1.3 Experimental design and procedure

Participants sat in front of a monitor (ASUS VG27AQA1A, 2560×1440, 100Hz), at a viewing distance of about 80 cm. The participants wore a data-glove on their real right hand (RH), which was hidden from view. The movements of the data glove were fed to a virtual hand (VH) presented on screen, behind the target dot, using Unity (Fig. 1A-C). Participants had to perform continuous grasping movements paced by a target dot presented centrally on screen, whose size continuously de- and increased (i.e., the dot’s diameter varied between 0-5 cm on screen, corresponding to about 3.6 degrees visual angle change in diameter) rhythmically at a predefined frequency (see below). Participants had to maintain fixation on the target dot and adjust their RH grasping movements, so that the VH grasping movements would follow/track the oscillation of the target dot, while a delay was consistently introduced between the RH and VH movements, according to the experimental condition (Fig1. A&B).

**Figure 1.**
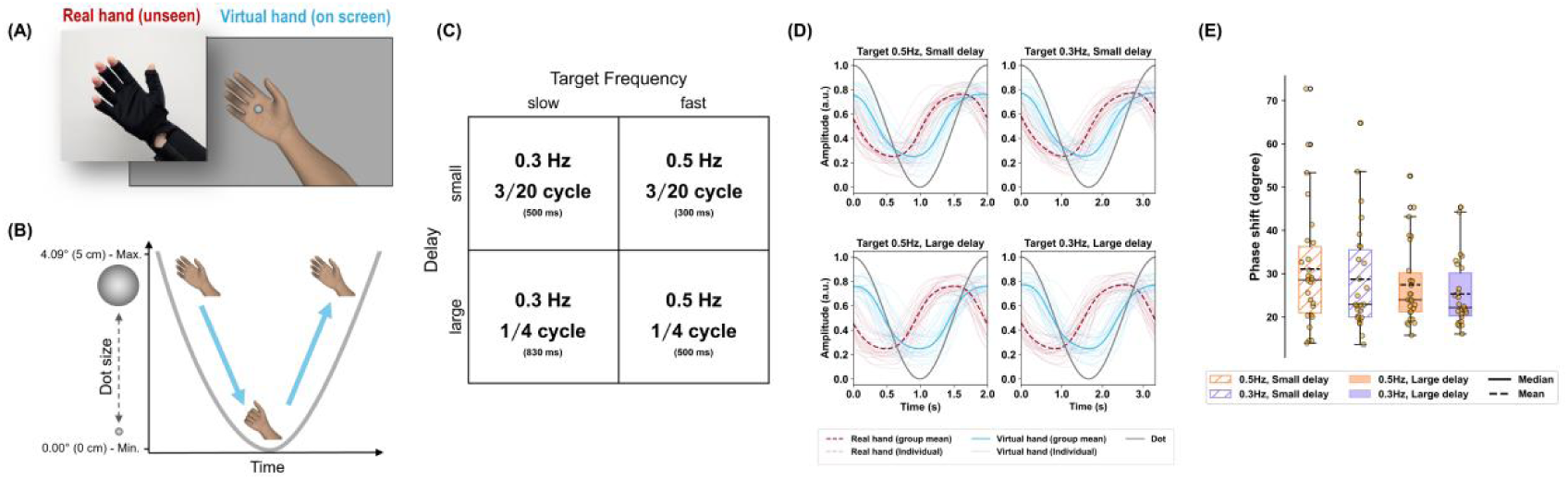
Experimental setup and design. **(A)** Participants wore a data glove on their real right hand (RH), which was hidden from view, while controlling a virtual hand (VH) displayed on the screen. The movements of the virtual hand were permanently delayed with respect to the actual movement; i.e., they lagged the participants’ movements by a certain time delay. **(B)** A target and fixation dot was presented centrally on screen, whose size continuously decreased and increased at a predefined frequency (see D) between 4.09-0 degrees visual angle. Participants were instructed to perform continuous grasping movements so that the VH followed the rhythmic size change of the dot, closing the VH as the dot decreased in size and reopening as it increased. As the VH movements were delayed, participants had to shift their actually executed hand movements in time to compensate for the delay. Each trial contained 18 continuous grasping cycles. **(C)** We varied the target (grasping) frequency (slow i.e. 0.3 Hz, or fast i.e. 0.5 Hz) and the amount of delay added to the VH movements (small delay i.e. 3/20^th^ of a cycle, or large delay i.e. 1/4^th^ of a cycle). This resulted in a 2 × 2 within-subject design with factors target frequency (slow vs. fast) and visuomotor delay (small vs. large). **(D)** RH and VH movements in each condition. The grey curve shows the target oscillation, the thick blue curve shows the grand-averaged VH movements, and the thick red curve shows the grand-averaged RH movements; the individual participants’ averaged movements are shown as thin lines. Note that, as instructed, in each condition, participants counteracted the visuomotor delay i.e. by phase-shifting their real hand movements to align the VH with the target oscillation. See Results for details. **(E)** Performance quantified by the phase shift between the virtual hand and the target for experimental condition. The boxplots show the median, mean, and inter-quartile range (IQR, whiskers extending to 1.5×IQR) with jittered dots representing individual value of participants. The performance was significantly modulated only by the delay level (*F(1,75)=6.49, p=0.0129; = −0.33*, linear mixed model), with larger delay level associated with better performance (i.e., smaller phase shift).

The experiment used a 2 x 2 within-subject factorial design (Fig. 1D), with two independent variables, *target frequency* and *delay* between RH and VH. Each factor had two levels: *target frequency* (*slow* 0.3 Hz or *fast* 0.5 Hz); *delay* (*big:* phase shift with 1/4 cycle or *small:* phase shift with 3/20 cycle, one cycle = one full grasp from open to close to open). It should be noted that we used fixed phase shifts instead of fixed time delays, to ensure the comparable grasp phase alignment with the beta band power dynamics reported in our previous study (Wang & Limanowski, 2023). In this case, for the slow target (0.3 Hz), the temporal delay was 0.833 s and 0.5 s, and for the fast target (0.5 Hz) 0.5 s and 0.3 s.

Prior to the actual experiment, the potential participants completed a training session, designed to assess whether participants could successfully perform the task; i.e., align the VH with the target oscillation. Participants who demonstrated sufficient task performance (averaged abs. phase shift error across the training session less than 60 degree; more details for the computation of absolute phase shift error (see Section 2.4.1) were subsequently invited to take part in the second EEG recording session, during which they performed the same task while EEG data was recorded. The interval between the training and EEG sessions was kept within 7 days.

At the beginning of the EEG recording session, participants were again introduced to the task and practiced each experimental condition. Once participants reported feeling confident in performing the task, the experimental session began. From this point onward, data collection—including hand movement, eye tracking, and EEG—was initiated.

In the EEG recording session, participants completed five blocks of 10.6 mins each. Each block contained 12 trials presented in random order, with each trial consisting of 18 continuous grasping movements under the same experimental condition. Therefore, each of the four experimental conditions was repeated three times per block, and no same condition appeared in two adjacent trials.

Each trial was separated by a 5 second fixation period, during which only a static target dot was presented. Participants were instructed to focus on the target dot and keep their hand still until the virtual hand appeared on the screen. This fixation period served as the baseline condition for the later EEG analysis. The upcoming condition was shortly announced by a text above the static target dot for the first 2 seconds of a trial, e.g. SLOW/SMALL, indicating the upcoming condition was slow target frequency and small delay. After completion of a block, participants were allowed to take a short break.

### 2.2 Data acquisition

#### Hand movements

Grasping movements were recorded using a data glove (5DT Data Glove) worn on the participant’s right hand. The glove measured finger flexion at a sampling rate of 100 Hz (through Unity), which was stored for later analysis and simultaneously mapped onto the fingers of a virtual hand model. In the virtual hand model, the thumb was fixed, and the flexion values of the remaining four fingers were averaged. In this way, participants controlled the finger movements of the virtual hand in real time—although a temporal delay was experimentally introduced (see Section 2.1.2, Experimental Design and Procedure).

For each participant, the data glove was calibrated after task instruction and prior to the practice movements. The flexion signal was calibrated individually such that a fully open hand corresponded to 0 and a fully closed hand to 1. This normalization was fixed for the remainder of the recording session. For analysis and visualization purposes, the values were inverted—so that 1 indicates a fully open hand and 0 a fully closed hand. Throughout the entire task, the measured RH movement values remained within the normalized range (0 < RH < 1; see Fig.S1).

#### EEG

EEG recordings were conducted using a 64-channel EEG system (actiCHamp Plus 64, Brain Products) with active, gel-based electrodes (actiCap Snap, Brain Products), mounting on an electrode cap (64 electrodes plus one ground electrode, Easycap, Brain Products) and trigger box (TriggerBox Plus, Brain Products). Data was recorded at a sampling rate of 1000 Hz. The ground electrode was placed on the forehead, and the online recording reference electrode was Cz. During the EEG recording session, trigger signals were sent from the experimental computer (running Unity) via a trigger box to synchronize the task events with the EEG data stream on the recording computer.

#### Eye tracking

Eye-gaze data was recorded using an eye tracker (GP3 HD eye tracker; Gazepoint), placed below the monitor, at a sampling rate of 60 Hz.

### 2.3 Data processing and analysis

#### 2.3.1 Behavioral performance

The participants’ performance accuracy (Fig.1E) was computed as the absolute phase shift between the target dot and the VH. This measure reflects how accurately participants performed the task, namely, to overcome the visuomotor conflict so that the VH closely tracked the rhythmic size changes of the target dot (see section 2.1.3 for details). The recorded averaged finger flexion (except thumb) represented the sinusoidal grasping movement (Fig. 1D).

First, the VH movement data was separated into four condition-specific epochs, each containing 15 trials with 18 continuous grasping cycles. Importantly, the grasp-cycle time windows used throughout this paper were defined according to **the** theoretical “perfect” grasp cycle, which is strictly aligned to the target dot’s size oscillation (max-min-max sequence). Thus, one grasping cycle corresponds to a complete open–close–open sequence as defined by the target trajectory (Fig. 1D). This cycle definition is based on the target dynamics rather than the actual movements performed by each participant. As participants were instructed to track the target, their executed grasping cycles were well aligned with the theoretical target-defined cycles. The first grasp of each trial was discarded from behavioral analyses, yielding 255 VH grasping movements per condition. Each single VH grasping was segmented using the time window that aligned with the corresponding one-oscillation period of the target dot (Fig. 1E). Note that further on, a single grasp cycle refers to this time window. The magnitude of the target dot was normalized between 0 (fully disappeared) and 1 (maximum size).

To estimate the averaged absolute phase shift error for each VH grasping, a Hilbert-transformation was applied to both the VH trajectory, and the corresponding target dot trajectory. The absolute phase difference was computed at each time point across a single target dot oscillation period, and then averaged across all 255 grasping movements per condition. Furthermore, we tested for potential differences in hand movement amplitude across experimental conditions (Fig. S1). For each participant, we computed the movement amplitude as the difference between its maximum and minimum measured finger flexion value in each individual grasping cycle. These values were then averaged across all 255 grasping cycles for each participant and each condition.

We also examined the continuity of tracking performance through the phase locking between the target dot movement and the VH movement through intersize phase clustering (ISPC). ISPC measures the consistency of the phase relationship between two trajectories across time regardless of the phase shift error (Cohen, 2014). Higher ISPC values indicate more continuous and stable tracking, with a value of 1 reflecting perfect phase locking.

#### 2.3.2 EEG pre-processing

EEG raw data was pre-processed in Matlab using EEGlab (Delorme & Makeig, 2004), along with several extensions (see below) and a customized pipeline, to perform resampling, line noise removal, filtering, bad channel detection and interpolation, independent components analysis (ICA), artifact rejection, and epoching. We down-sampled the data to 300 Hz, used zapline-plus (de Cheveigné, 2020) to remove 50 Hz line noise, and applied a 1 Hz high-pass filter to improve ICA performance. Before ICA, the potential bad channels were identified with clean_rawdata (Kothe & Makeig, 2013, Delorme & Makeig, 2004) and interpolated. ICA (PICARD (Ablin et al., 2018)) was performed together with ICLabel (artifact threshold 0.9; (Pion-Tonachini et al., 2019)) to detect bad ICs (ocular, and muscular). Since we targeted low frequencies (e.g., 0.3 Hz), we copied the ICA weights back to the unfiltered data (after down-sampling and line noise removal) and rejected the bad ICs afterwards. Visually inspected, the removed ICs were very similar for each individual across the 5 blocks. Before epoching, EEG data was high-pass filtered at 0.25 Hz. No single-trial rejection was applied. Artifact-free EEG data was then segmented into five epochs: four condition-specific epochs, each corresponding to one experimental condition and containing 15 trials (each trial including 18 VH grasping movements), and one baseline epoch comprising 60 trials of the 5-second fixation period. We refer to these condition-specific epochs as the pre-processed long-epoch data.

#### 2.3.3 EEG power spectral density (PSD)

The PSD for each experimental condition was computed with Welch‘s method using the Fieldtrip toolbox (Oostenveld et al., 2011) in Matlab. The condition-specific pre-processed long-epoch data was further segmented into trials consisting of four concatenated grasping cycles, with an overlap of three cycles, yielding 255 trials per condition. For each trial, a Hanning window was applied, and PSDs were estimated from 0.25 Hz to 90 Hz, zero-padding to 20s (frequency resolution of 0.5 Hz). Power values were converted into decibels (dB). The same method was applied to the 5s-fixation period data (baseline), but without overlapping between trials. Finally, for each channel, the trial-averaged PSD was baseline corrected by subtracting the trial-averaged baseline PSD.

In addition to the first harmonic power, we also estimated the theta band power (4-7 Hz), alpha band power (8-13 Hz), beta band power (14-30 Hz) and gamma band power (31-70 Hz) by averaging the spectral power across these frequency bins.

#### 2.3.4 Granger prediction between EEG and hand movement trajectories

We performed a spectral Granger prediction analysis (Barnett & Seth, 2015) to examine the directionality of interactions between EEG activity and hand movement trajectories, including both the real hand (RH) and virtual hand (VH). The analysis focused on the CPz-centered cluster (CPz, CP1, CP2), which served as our region of interest (ROI). This cluster was selected because its f1 response power showed a reliable modulation by the delay (Fig. 2C). We examined both directions of information flow: (i) whether, on average, hand movements predicted EEG activity (RH/VH → EEG) or (ii) EEG activity predicted subsequent hand movements (EEG → RH/VH).

**Figure 2.**
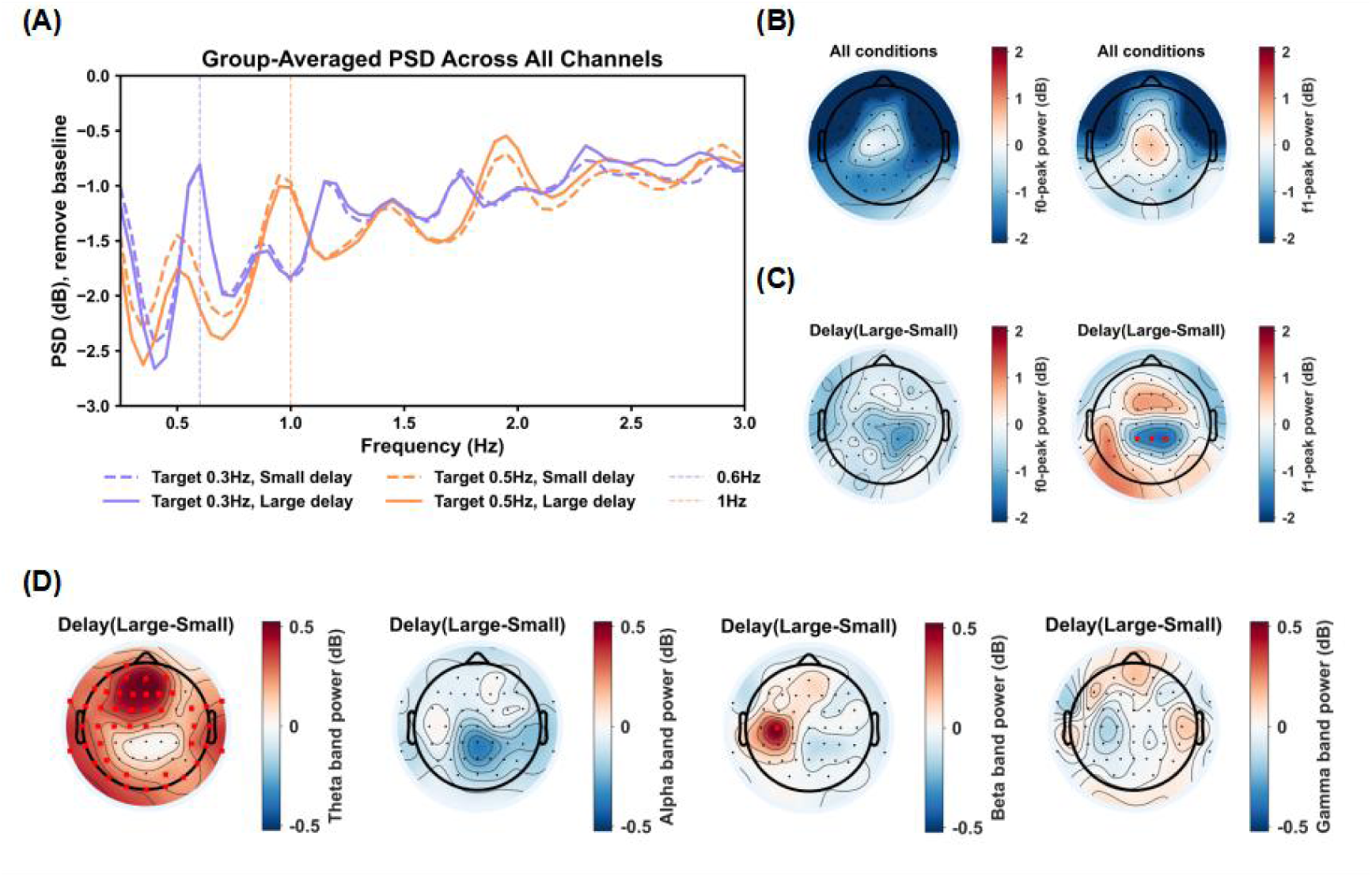
Visuomotor delay modulated f0, f1-components as well as other frequency bands. **A:** Group-averaged, baseline corrected power spectral densities (PSD) across all 64 EEG channels from 0.3 to 3Hz for each experimental condition. EEG power showed strong peaks at the first harmonic of the key movement frequency; i.e., conditions with a target frequency of 0.5Hz showed a prominent 1 Hz peak in PSDs (red lines), whereas conditions with a target frequency of 0.3 Hz showed a prominent 0.6 Hz peak in PSDs (blue lines). **B:** Power topology at the target frequency (f0; left) as well as at its first harmonic component (f1), averaged across all experimental conditions. **C:** Power topology of the main effect delay for the f0-peak (left) as well as f1-peak (right); i.e., the difference between large and small delay conditions. Red dots mark sensors showing a significant main effect of delay (FDR corrected p<0.05). **D:** Spectral EEG power topologies of the main effect of delay at theta (4-7 Hz), alpha (8-12 Hz), beta (13-30 Hz) and gamma (31-70 Hz) frequency bands. Red dots mark sensors showing a significant main effect of delay (FDR corrected p<0.05).

Although the VH was a delayed version of the RH, the analysis time windows were aligned to the oscillation period of the target dot (or grasping cycle), yielding meaningful phase differences between the RH and VH time series (see Supplementary Figs. S6-S7). Both EEG and hand movement data was extracted from the condition-specific, pre-processed long-epoched data (containing 18 grasping cycles per trial). Note that since RH and VH were evaluated in separate Granger models, the results reflect directed coupling within each EEG – hand movement pair rather than unique contributions of one hand signal conditioned on the other.

For the main analysis, the long continuous epochs (18-cycles per trial) were re-epoched into 165 new trials per condition. Specifically, we discarded the first grasping cycle from the original 18-cycle sequences and then generated shorter epochs comprising consecutive seven cycles, with each new epoch overlapping by six cycles with the preceding one. To match the sampling rate of the EEG data (300 Hz), the hand movement data was up-sampled to 300 Hz.

Spectral Granger connectivity was computed using a multivariate state-space Granger approach (Barnett & Seth, 2015) implemented in the MNE-Connectivity toolbox (spectral_connectivity_time(); (Binns et al., 2026)). We tested two specific prediction directions of interest: stimulus-driven entrainment (RH/VH → EEG) and internal prediction (EEG → RH/VH). For each condition, we computed spectral Granger values in both directions and quantified the dominant direction of information flow as the difference between EEG → hand and hand → EEG Granger values (Fig. 3A).

**Figure 3.**
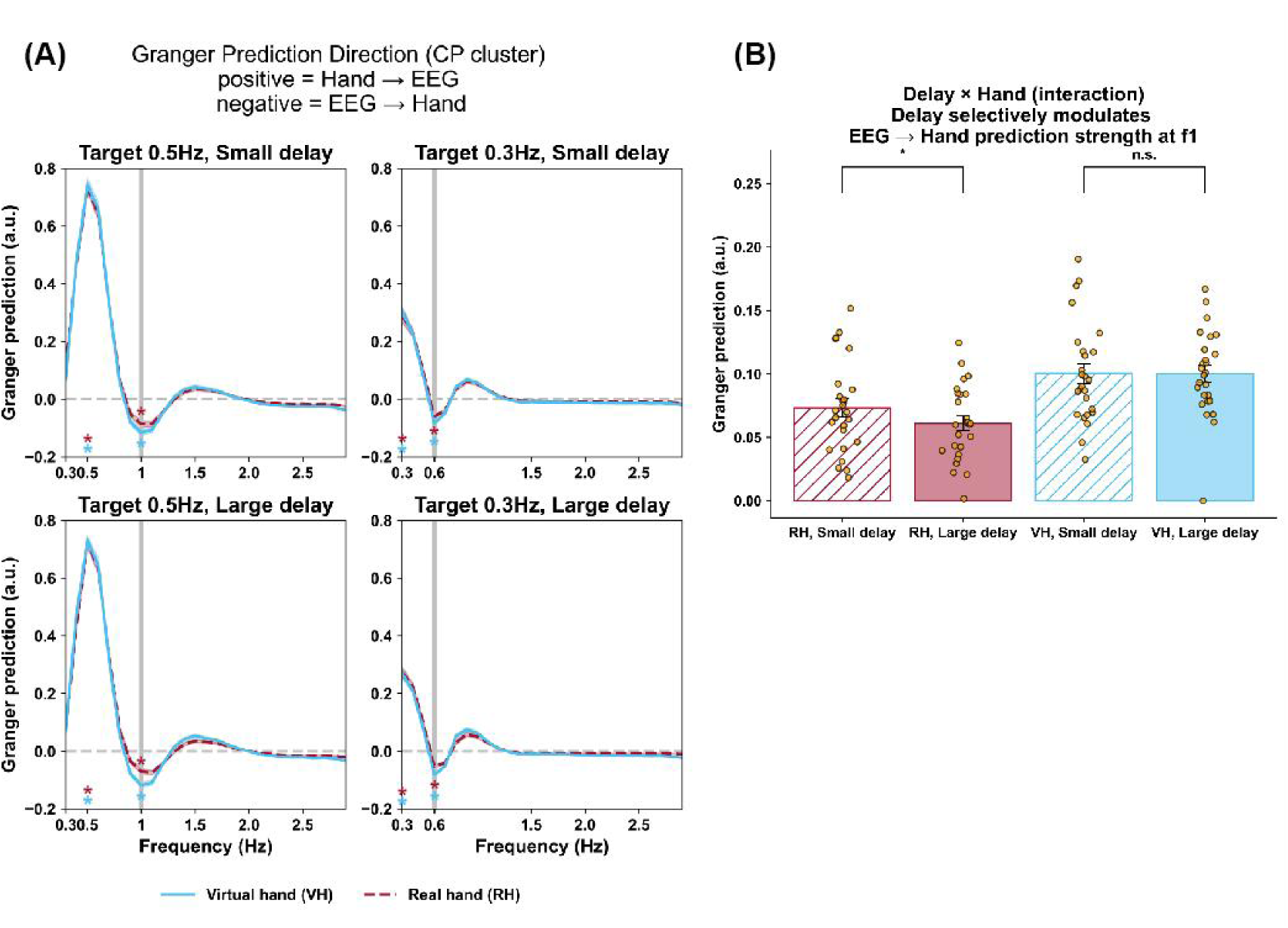
Spectral Granger analysis of EEG vs real and virtual hand movements at CP cluster electrodes. **(A)** Spectral Granger prediction among EEG data (CPz, Cp1 and Cp2), real hand (RH), and virtual hand (VH) movement trajectories, across all four experimental conditions. Dominant Granger influence direction was estimated through the difference between the spectral granger prediction from hand (RH or VH) to EEG and the spectral granger prediction from EEG to hand (from 0.3-3 Hz in 0.1 Hz increments). The positive peaks at 0.3 and 0.5 Hz, the target/hand movement frequency (f0), indicate that the dominant influence direction was biased from hand to EEG: VH/RH→EEG. Conversely, the negative peaks at 0.6 and 1 Hz indicate that the dominant influence direction was biased from EEG to hand: EEG→VH/RH, aligned with the first harmonic frequency f1 of the target/hand movement frequencies. Statistical tests were applied on both f0 and f1. Asterisks indicate peaks significantly different to zero for the VH (blue curves) or RH conditions (red curves, FDR corrected *p*<0.05) at the group level. **(B)** Group means of absolute Granger prediction strengths (Granger prediction bias towards EEG→RH) at f1 frequency (averaged across target frequencies). Error bars represent standard errors of the mean. Dots represent individual participant values. The prediction strength EEG→RH was significantly modulated by the delay condition, with stronger prediction bias towards EEG→RH under small than under large delays (*t(25)=*2.40*, p*= 0.024), wherever there was no difference in the EEG→VH predictions at small vs large delays (*t(25)=*0.041*, p=*0.96). See Results for details.

Before the main analysis, to ensure that reliable Granger-prediction effects were observable in the data, we performed a permutation-based control analysis to confirm that the observed directional effects were not due to chance. This procedure was applied only for quick validation (for details, see Supplementary Information *Granger prediction*).

The Granger-prediction analysis was implemented with a fixed prediction window of 20 samples (N = 20 at 300 Hz, ≈ 67 ms). Shorter history windows of N = 5 (≈ 17 ms) did not yield reliable estimates of the directional Granger prediction from EEG to Hand at the f1 frequency (for target 0.3 Hz condition) in the permutation control test. (see Supplementary Information *Granger prediction*).

#### 2.3.5 Phase-amplitude coupling on local and cross-site electrodes

Phase-amplitude coupling (PAC) is a method to quantitatively describe the phenomena that the amplitude of high-frequency oscillations is modulated by the phase of low-frequency rhythms (Canolty et al.,2010; Lakatos et al. 2008; Tort et al. 2010). In our previous visuomotor conflict study (Wang & Limanowski 2023), we reported an increased PAC between the phase of the first harmonic of the grasping rhythm (e.g. 1 Hz) and Beta (14-28 Hz) power amplitude. Following up on the detection of the significant power modulation of the first harmonic response (f1), theta, as well as beta oscillations under the large delay condition in PSD (Fig. 2), we asked whether the beta amplitude (at electrode C3) was modulated by the phase of either of these lower-frequency oscillations. Specifically, we tested local PACs as well as the following cross-site PACs: C3(f1)-C3(beta), C3(theta)-C3(beta), CPz(f1)-C3(beta), Fz(f1)-C3(beta).

We computed PAC using the modulation index method (MI; Tort et al. 2010) implemented in tensorpac (Combrisson et al., 2020) as follows: we computed the time-frequency representations (TFRs) for the low frequencies (f1 of 1 Hz, 0.6Hz, as well as theta band 4-7 Hz) and the high frequencies (beta band (13-30 Hz), for each trial (including 18 continuous target oscillation cycles) and each condition, using Morlet wavelets with six frequency cycles. The estimated phase and amplitude were then concatenated along the time axis for each condition and we computed the MI of the PAC by examining the amplitude distribution over 18 phase bins (i.e. 20 degrees per phase bin). We used a permutation approach to estimate the z value from distribution of the surrogated MI of the PAC, by swapping amplitudes time blocks 200 times (Bahramisharif et al., 2013).

We selected MI over other alternative PAC metrics because it is insensitive to amplitude/power magnitude (Tort et al. 2010). As the visuomotor delay significantly modulated the beta band as well as the theta band power, by using an amplitude-invariant measure, we ensured that observed variations in PAC reflect changes in cross-frequency coupling rather than trivial differences in power levels.

#### 2.4.5 Eye gaze points

Since participants were instructed to maintain fixation on the central fixation point during the task, we tested whether gaze fixation was comparable across all conditions. Gaze positions were recorded in normalized screen coordinates (range 0–1), with (0.5, 0.5) corresponding to the center of the display, which was spatially aligned with the fixation point and the middle of the target dot. For each gaze sample, the Euclidean distance to the fixation point at the screen center (0.5, 0.5) was computed. The mean gaze-to-center distance for each condition was obtained by first averaging the distances across all samples within each trial and then averaging across the four trials of the same condition. All gaze points were classified online during recording as either valid or invalid automatically by the eye-tracking system.. An invalid sample means that participant’s eyes could not be reliably detected from the eye-tracking device. A valid ratio of gaze points was calculated separately for each of the four experimental conditions. Participants were included in the gaze data analyses only if the valid ratio exceeded 90% in all conditions. Based on this criterion, 18 of the 26 participants were available for further analyses.

### 2.4 Statistical analysis

#### 2.4.1 Behavioral performance

A linear mixed-effects model (Fig. 2C) was conducted for the within-subject 2×2 factorial design with the factors target frequency (0.5 Hz, 0.3 Hz) and delay (small, large):

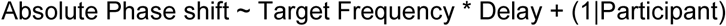

Absolute phase shift was introduced as the dependent variable, while target frequency and delay were included as fixed effects along with their interaction. A random intercept per participant accounted for within-subject dependencies. Fixed-effect significance was examined using linear mixed-effects models implemented in MATLAB (*fitlme()*). Models were fitted with restricted maximum likelihood (REML) estimation and effects coding for categorical predictors. We obtained F-statistics and p-values for each fixed effect using Type III tests with Satterthwaite-approximated denominator degrees of freedom, ensuring results are directly comparable to traditional repeated-measures ANOVA while appropriately accounting for the random subject intercept. Model residuals were inspected for normality, and no violations were detected. Within-subject effect sizes (Cohen’s dz) were computed post-hoc from participant-level contrast scores derived from condition means. Main-effect contrasts were defined as large minus small for the delay factor and 0.5-Hz minus 0.3-Hz for the target-frequency factor. The interaction effect was quantified as the difference between these delay effects across target frequencies (i.e., (large − small) at 0.5 Hz − (large − small) at 0.3 Hz). We conducted an analogous analysis to test for differences in movement amplitude across conditions (see Fig. S1) as well as for the gaze points (see Table S7).

#### 2.4.2 EEG PSD and topology

To address whether the peak power at the target frequency (f0), or its first harmonic (f1), was modulated by experimental conditions, we calculated a linear mixed model for f0 and f1, respectively, within a 2×2 within-subject factional design: frequency (0.3/0.6Hz, 0.5/1Hz) x delay(small, large):

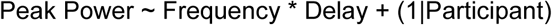

The linear mixed model was computed separately for each EEG channel. The implementation of the linear mixed-effects modeling was the same as described above and applied to the behavioral data (see Section 2.5.1). Model residuals were inspected for normality, and no violations were detected. Multiple comparison was corrected using the Benjamini-Hochberg procedure to adjust the false discovery rate (FDR). All multiple comparison corrections were performed using the same procedure. Effect sizes were estimated post-hoc through Cohen’s d_z_. Statistical tests using linear mixed model were implemented through customized Matlab scripts.

#### 2.4.3 Granger prediction between EEG and hand movement data

For each participant and experimental condition, we extracted the directional Granger prediction values at the first harmonic of the target frequency: 1 Hz for the 0.5 Hz target and 0.6 Hz for the 0.3 Hz target, separately for RH and VH (Fig. 3A). First, one-sample t-tests were performed to test whether EEG → hand prediction values at the first harmonic were significantly different from zero across participants, separately for each condition and hand modality. Multiple comparisons were corrected using FDR. To examine how experimental conditions modulated predictive EEG activity, we conducted a three-way repeated-measures ANOVA with the within-subject factors: hand modality (RH, VH), target frequency (0.3 Hz, 0.5 Hz), and delay (small, large). The dependent variable was the absolute value of directional Granger prediction value at the f1 frequency. Where significant interactions were identified, paired-sample t-tests were used as post hoc comparisons. Statistical tests were implemented through customized python script using *scipy* and *statsmodel*.

#### 2.4.4 Phase-amplitude coupling on local and cross-site electrodes

At the single subject level, the PAC was z-normalized; the z-score indicated the observed PAC strength relative to a surrogate distribution mean. I.e., PAC was normalized to a z-score by comparing the observed MI with a surrogate distribution generated by swapping amplitude time blocks 200 times. The mean of this surrogate distribution corresponds to the average PAC expected when the temporal relationship between phase and amplitude is disrupted, thereby providing an estimate of chance-level coupling. At the group level, we tested whether PAC exists in our visuomotor conflict conditions by computing the mean PAC z-scores averaging across all conditions for each participant and performing a one-sample two-sided t-test against zero. A significant greater-than-zero result indicates that PAC is greater than null expectation across subjects.

To address whether PAC was modulated by experimental conditions, we performed a linear mixed model within a 2×2 within-subject factional design: target frequency (0.3Hz, 0.5Hz) x delay(small, large), similarly implemented like described in session 2.5.1 and 2.5.2 for each 37 combinations of PAC:

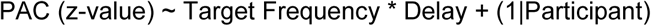

We corrected for multiple comparisons using the false discovery rate (FDR) method with the Benjamini–Hochberg procedure. Effect sizes were estimated post-hoc through Cohen’s d_z_. Statistical tests using linear mixed model were implemented through customized Matlab script.

## 3. Results

### 3.1 Behavioral results

Participants were able to successfully perform the rhythmic grasping task; i.e., to counteract the visual feedback delay and align the VH with the target oscillation under all experimental conditions. Fig. 1D illustrates the averaged hand trajectories during the full grasping cycle. Overall, tracking performance, movement amplitude, and movement phase consistency was comparable across experimental condition (Fig. 1E; cf. Figs. S1-S2 for comparisons of movement amplitudes and phase consistencies). Tracking was slightly better at large than small delays (i.e., participants made larger target tracking errors under small delays, *F(1,75)=6.49, p=0.0129;* dz *= −0.33*), while movement phase consistency was somewhat lower at large than small delays (*F(1,75)=14.43, p=0.00029;* dz *= −0.67*).

### 3.2 Spectral EEG effects related to visuomotor delay magnitude

The averaged PSD across all 64 EEG channels revealed clear f0- as well as f1- peaks corresponding to the target frequencies (Fig. 2A); i.e., prominent 0.5/1 Hz peaks in the 0.5 Hz conditions and 0.3/0.6 Hz peak in the 0.3 Hz conditions. The f0- as well as the f1-component shared a similar topology during each task compared against a baseline condition without target and hand movement, with higher power at central electrodes (Fig. 2B, cf. Fig. S4). Furthermore, a linear mixed-effects model of f1- peak power at each electrode revealed a significant main effect of delay at the post-central midline electrodes CPz (*F(1, 75)*=14.02, *p*=0.00035, FDR corr. *p*=0.011, dz=-0.84), CP1(*F(1, 75)*=11.72, *p*=0.0010, FDR corr. *p*=0.022, dz=-0.80), and CP2(*F(1, 75)*=14.83, *p*=0.00025, FDR corr. *p*=0.011, dz=-0.92; Fig. 2C). We will refer to these three electrodes (CPz, CP1, CP2) as the “CP cluster”. At these electrodes, power at the respective harmonic target frequency (f1) was significantly attenuated during large compared to small delays (Fig. 3C; cf. Fig. S5 for the main effect of target frequency). There was a similar, but non-significant attenuation of f0-peak power around post-central right side (CPz, CP2) during large compared with small delay.

We further tested for delay related differences in theta (4-7 Hz), alpha (8-12 Hz), beta (13-30 Hz), and gamma (31-70 Hz) band neural oscillation power. This analysis revealed a significant main effect of delay in the theta frequency range, with the peak effect located at mid-frontal electrodes (Fig. 2D); theta power was elevated during large > small delays. Beta power (13-30 Hz) at electrode C3, likely corresponding to contralateral sensorimotor cortex, significantly increased during movements under large > small delays (*F(1, 75)*=12.93, *p*=0.00058, FDR corr. *p*=0.037, dz=0.54). Alpha power was attenuated at posterior electrodes (CPz, Pz; Fig.2D) during large compared with small delays, but this effect did not reach significance. Gamma power was relatively attenuated at electrode C3 during large compared with small delays, but this effect did not reach significance. See Fig. 2D for details. The main effect of target frequency see Fig. S5.

### 3.3 Spectral Granger prediction analysis between EEG and hand movement trajectories

The spectral Granger prediction analysis between EEG activity at the CP cluster and RH and VH movement revealed, across all conditions, positive peaks at the target frequencies (f0 i.e., 0.3/0.5 Hz), but negative peaks at the harmonic of the respective target frequency, for both VH and RH signals (Fig. 3A and Table S3&S4). The positive peaks indicate that VH/RH signals predicted EEG (CP cluster) activity at the respective target frequency, more strongly than vice versa. Conversely, the negative peaks indicate that EEG activity at the CP cluster more strongly predicted upcoming movement transitions (e.g., from the hand closing to opening and vice versa), which occurred twice per movement cycle.

Next, we tested for conditional differences between these Granger prediction biases. At the target frequencies f0, there was a significant main effect of target frequency (*F(1, 25)*=1288, *p*<0.05, 0.5 Hz > 0.3 Hz), and a significant main effect of delay (*F(1, 25)*=6.21, *p*<0.05, large delay < small delay; cf. Table S6). No other effects were significant.

At the f1-harmonics, there was a significant main effect of hand modality (*F(1, 25)*=84.37, *p*<0.05), with a larger predictive bias (i.e., absolute value) for EEG→VH than EEG→RH. There was no significant effect of delay, but a significant main effect of target frequency f0 (*F(1, 25)*=16.64, *p*<0.05, 0.5 Hz>0.3 Hz) and, crucially, a significant interaction between delay and hand (*F(1, 25)*=6.45, *p*=0.018; cf. Table S5). To follow up on this interaction effect, we calculated post-hoc paired t-tests contrasting the Granger prediction direction between RH small vs large delay, and VH small vs large delay conditions, respectively, pooling across target frequencies in each comparison. This revealed that the EEG→RH prediction was significantly modulated by delay, with stronger prediction values under small than under large delays (*t(25)=*2.40*, p*=0.024). In contrast, this difference was non-significant for the EEG→VH prediction (*t(25)*=0.041, *p*=0.96).

### 3.4 Phase-amplitude coupling

Our PAC analysis focused on the potential local and cross-site coupling between oscillations showing delay effects at f1 or theta frequencies (potentially constituting top-down “control signals”, see Introduction) and beta oscillations at C3. This analysis (Fig. 4) indicated several nontrivial local and cross-site couplings between the phase at f1 (1 Hz/0.6 Hz) and theta band (4-7 Hz) phase and amplitude at beta band (13-30 Hz). All of those PAC pairs were significantly above the surrogate baseline threshold; i.e., (C3(f1)-C3(beta): *t(27)*=7.07, FDR corr. *p*=8.27e-7; C3(theta)-C3(beta): *t(27)*=2.74, FDR corr. *p*=0.011; CPz(f1)-C3(beta): *t(27)*=4.57, FDR corr. *p*=0.00023; Fz(theta)-C3(beta): *t(27)*=3.06, FDR corr. *p*=0.0070; Fig. 4A).

**Figure 4.**
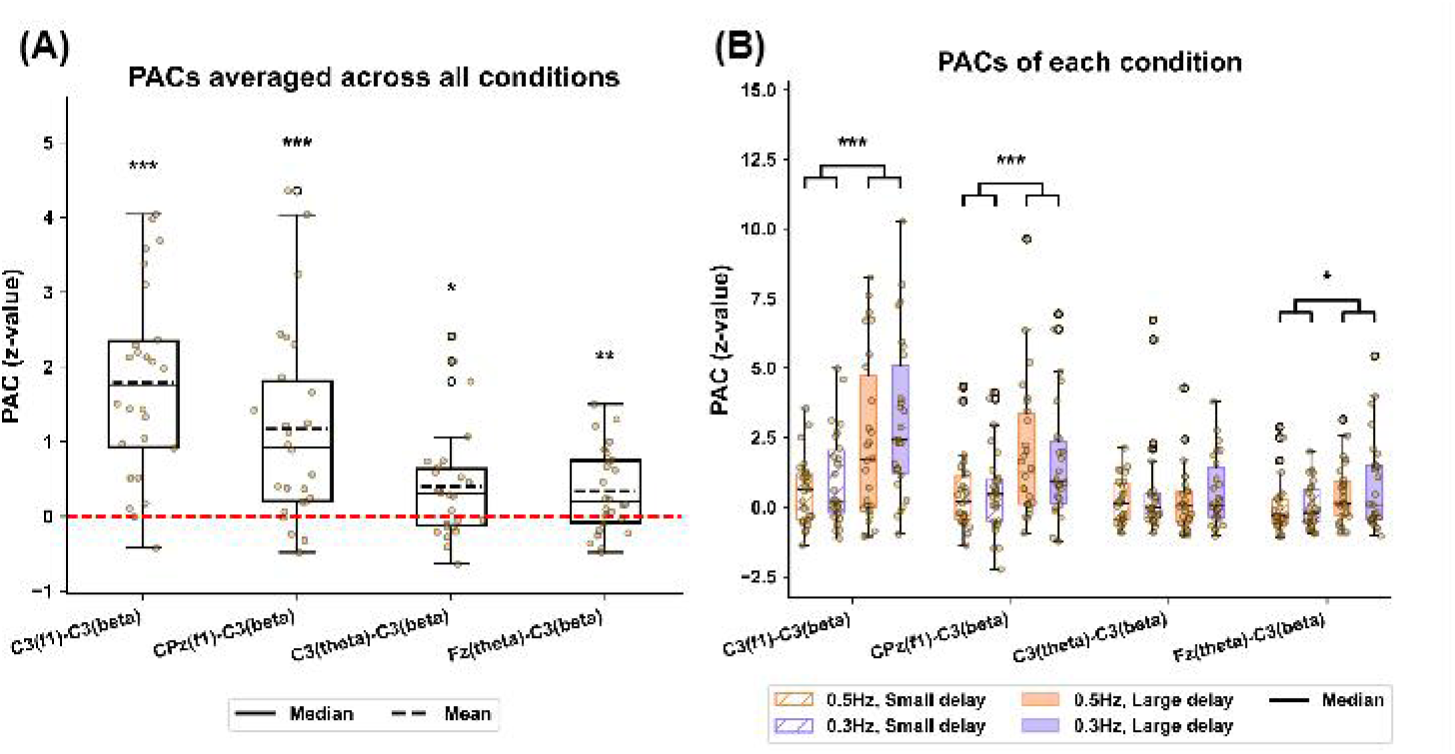
Phase-amplitude coupling. **(A)** Boxplots of the mean PAC values (averaged across all conditions) for f1-beta and theta-beta couplings, locally at C3 and cross-sites (CPz-C3, Fz-C3). The mean PACs were tested by one-sample two-tailed t-test against the surrogate baseline (z=0, red dashed line). All estimated PACs at group level exceeded this chosen threshold (FDR corr. *ps*<0.05) **(B)** Boxplots of the PAC values per condition. The f1-beta PACs were significantly modulated by delay (Large > small; C3-C3: FDR corr. *p*<0.001; Cpz-C3: FDR corr. *p*<0.001). The theta-beta PAC across Fz-C3 was also significantly modulated by delay (Large > small; FDR corr. *p*<0.05), while theta-beta locally at C3-C3 was not (FDR corr. *p*=0.93). For the boxplots, each dot represents an individual participant. The box indicates the interquartile range (25%-75%), the solid horizontal line denotes the median, and the dashed line indicates the mean. Data beyond 1.5 × the interquartile range (IQR) is indicated with empty circles. * corr. *p*<0.05, ** corr. *p*<0.01,*** corr. *p*<0.001, FDR-corrected.

Furthermore, there were significant effects of delay on f1-beta PAC at site C3-C3 (*F(1,75)*=20.83, FDR corr. *p*=0.00023, dz=0.73) and cross-sites CPz-C3 (*F(1,75)=*12.53, FDR corr. *p*=0.0042, dz=0.54). For theta-beta PAC we only found a significant effect at cross-site Fz-C3 (*F(1,75)*=7.10, FDR corr. *p* =0.036, dz =0.43), not at C3 (*F(1,75)*=0.0083, FDR corr. *p*=0.93, dz=-0.019). See Figure 4B.

## 4. Discussion

In our virtual reality-based hand-target phase matching task under delayed visual hand movement feedback, we identified several spectral EEG changes related to visuomotor control under visuo-proprioceptive conflict: (1) Oscillations “entrained” at the movement harmonics showed significant power modulations depending on the amount of visual feedback delay, and delay-dependent (Granger) predictive biases between EEG and kinematic signals; (2) oscillatory power in the beta and theta ranges was modulated by the amount of visual feedback delay; and (3) there were delay-dependent changes in the PAC among the identified key sensors. We now unpack these results in detail.

Firstly, EEG spectral activity over central-parietal electrodes showed clear peaks at key movement frequencies. This result suggests that synchronized neuronal activity, observed over central-parietal electrodes, reflects key variables related to movement kinematics i.e. action timing. The response peaking at the harmonics could mean a preferential “entrainment” of neural activity at the frequency of hand movement transitions (i.e., from closing to opening) instead of full cycles. As we have previously argued, in this kind of oscillatory movement task, the transitions between opening and closing are most informative for evaluating hand-target alignment (Wang & Limanowski, 2023; but entrainment at harmonics could also result from mere stimulus presentation at sub-delta frequencies, cf. Gomez-Ramirez et al., 2011).

Crucially, our Granger prediction analysis showed that, at these movement harmonic frequencies, EEG activity measured at the above central-parietal electrodes (CP cluster) predicted VH/RH movement signals, rather than the other way around. This could indicate a cortical prediction of visual and somatosensory (proprioceptive) signals relating to transitioning between the two different hand movements i.e., opening and closing. In principle, this speaks to the idea of state estimation in the posterior parietal cortex (PPC; Wolpert et al., 1998; Medendorp & Heed, 2019).

The predictive bias was stronger for VH than for RH signals, which could suggest a generally stronger prediction of visual than somatosensory signals at the movement transitions. Notably, both the entrained power and the Granger prediction bias were, at the central-parietal site, modulated by the amount of visuo-proprioceptive conflict: The “entrained” power at harmonic frequencies was lower under large than small delays (in line with a previous, but non-significant finding in Wang & Limanowski, 2023). The Granger predictive bias was likewise lower under large than small delays—but only for the prediction of RH signals, not VH (visual) signals. One potential interpretation of the above effects is that the central-parietal site relatively attenuated the prediction of RH movements under large delays (i.e., larger visuo-proprioceptive conflict).

Note that this could also mean that the PPC was mainly predicting visual movement transitions (of the VH and/or the target), and the prediction of RH signals was merely a by-product of their relative similarity to visual signals—which was lower under larger compared with small delays. These interpretations are not mutually exclusive, but future work with more specific study designs is needed to potentially disentangle them.

Secondly, beta power over electrode C3 increased significantly under large compared with small delays. While contralateral beta power attenuation is typically associated with motor activity (Salmelin & Hari, 1994), in our study, there was no effect of movement amplitude on these signals. Therefore, another interpretation of these signals is as somatosensory: Indeed, electrode C3 is thought to reflect signals from the primary somatosensory cortex (S1, Bernier et al., 2009; Kelly & Folger, 1999). Attenuation of beta (and alpha) oscillations recorded over sensorimotor areas has been frequently linked to somatosensory attention (Lebar et al., 2017; Bauer et al., 2012; van Ede et al., 2010, 2011; Keil et al., 2016; Jones et al., 2010). In line with this work, the relative increase in beta power under large compared with small delays would suggest relatively attenuated somatosensory attention. We observed a complementary pattern for posterior (likely, visual cortical) alpha oscillations; which were relatively augmented during large delays (but non-significantly so). This also aligns with previous reports in passive intersensory, visuo-tactile attention studies (Keil et al., 2014; Bauer et al., 2012; Lebar et al., 2017).

Together, our results suggest relatively attenuated somatosensory processing, and (somewhat) augmented visual processing, in the respective sensory cortices, during large compared with small visual feedback delays. This resonates with the hypothesis that visual, relative to somatosensory information, may be preferentially processed under conflict between visual and proprioceptive limb position estimates, to enable visually guided action under visuo-proprioceptive conflict (see Introduction), and supplements the notion of similarly biased sensory prediction in the parietal cortex (see above).

There are potential alternative interpretations of the observed C3 beta power modulation. Beta oscillations themselves have not only been implied in intersensory, but also in temporal attention (van Ede et al., 2011; Nobre et al., 2001; cf. Keil et al., 2016; Kulashekhar et al., 2016). Specifically, sensorimotor beta may reflect temporal predictions sent from the motor system to sensory regions for sensorimotor coordination (Morillon & Baillet, 2017; Fujioka et al., 2015; Kulashekhar et al., 2016; Pollok et al., 2008; Saleh et al., 2010; Arnal et al., 2014, 2015). Our task involved timing and predicting movement transitions, accompanied by an increase of sensorimotor beta in conditions where performance was overall better (large delays). It should be noted that movement phase consistency was lower under small delays, which could also hint at a more flexible adjustment of temporal predictions and attention in service of reducing spontaneous tracking errors. Furthermore, motor beta oscillations may be modulated by action observation (Press et al., 2011; Coll et al., 2017; Hari, 2006; He et al., 2018), including in scenarios requiring visuomotor self-other distinction and coordination (Bolt & Loehr, 2024). Thus, the C3 beta modulation by delays we observed may have reflected a modulation of motor cortical activity in response to action control under the differently delayed visual movements, instead of, or in addition to somatosensory attenuation. This interpretation is not mutually exclusive with the idea of somatosensory attenuation, but these ideas should be contrasted explicitly by future work.

Theta power was also significantly enhanced under large compared with small delays, most strongly so at mid-frontal electrodes. We have previously observed increased mid-frontal theta during visual hand-target matching under visuo-proprioceptive conflict relative to no-conflict (Limanowski et al., 2020). Here, theta power scaled with conflict. This aligns nicely with previous associations of mid-frontal theta power with attentional control in inter-sensory or visuomotor tasks (Watanabe et al., 2021; Murray et al., 2025; Spooner et al., 2019; Drew et al., 2026). Together, these results support the hypothesis that mid-frontal theta power indicates the need for top-down control (Ullsperger et al., 2014; Cavanagh and Frank 2014; Cohen and Donner 2013; Zavala et al., 2018). Specifically, our results suggest that theta scales proportionally with the amount of control needed in a visuomotor task.

It should be noted that participants matched the target significantly better with their grasping movements under large compared to small delays, although these differences were only small. Previous work has suggested that, under smaller visuo-proprioceptive conflicts, conflicting visual information may more strongly bias proprioceptive information through (Bayes-optimal) multisensory integration than under larger conflicts, where visual and proprioceptive signals may not be integrated (Botvinick & Cohen, 1998; Lloyd, 2007; Vigh & Limanowski, 2025). In our task, an analogous integration of conflicting (i.e., delayed) visual with somatosensory signals could have impaired hand-target matching with the VH, or distracted from using purely visual signals. Under larger conflicts, the conflicting visual and proprioceptive hand movement signals could, potentially, be more clearly separated. Importantly, the separation and conflict resolution (cf. Balslev et al., 2004; Bernier et al., 2009) could have been augmented by increased control and the adjustment of visual vs somatosensory attention; i.e., as indicated by changes in mid-frontal theta, and occipital vs sensorimotor alpha-/beta-oscillatory power.

Thirdly, beta power amplitude at electrode C3 coupled with the phase of oscillations at the harmonic movement frequency locally and at central-parietal sensors, and, somewhat weaker, also with the phase of mid-frontal theta. These couplings were enhanced under large delays. The modulation by the phase of key movement frequencies aligns with previous reports of “entrainment” by movement of sensorimotor beta oscillations (Miller et al., 2012). We have previously linked harmonic-beta PAC to rhythmic sensory attention (Wang & Limanowski, 2023); i.e., suggesting that in a periodic movement task like ours, sensory attention can be in-or decreased to align with behaviorally important time points, such as the transitions from closing to opening the hand. Our present findings suggest that somatosensory attention can be modulated in this way, potentially further augmenting the attenuation of distracting somatosensory information at particularly important time points during a visuomotor adaptation task. Furthermore, the coupling of theta and alpha/beta oscillations has been suggested as a potential mechanism conveying top-down control and response inhibition (Chacko et al., 2018; Watanabe et al., 2021; Wendiggensen et al., 2023). Thus, our results point to two potential underlying control signals modulating sensorimotor beta. Based on our spectral topography, these signals may come from the prefrontal cortex (e.g., through performance monitoring and the need for control by the anterior cingulate cortex, see Ullsperger et al., 2014) or the posterior parietal cortex (through bodily and environmental state estimation, see e.g., Medendorp & Heed, 2019).

Our results have to be interpreted in light of the following limitations. Firstly, while target frequency was not our primary factor of interest, we found significant target frequency effects in several of the investigated frequency bands (see Supplement). We speculate that this effect reflects differences in the window lengths chosen in the calculation of the PSD rather than purely physiological effects. I.e., the PSD analysis used the same number of movement cycles across frequencies; and longer windows (0.3 Hz) can increase estimated spectral power due to improved frequency resolution and reduced variance, which can produce sharper and more stable spectral peaks. Previous studies (Toma et al., 2022; Nordin et al., 2019) have reported decreased alpha- and beta band power during higher movement rate in sensorimotor regions, and while our findings are broadly consistent with this trend, the effects observed extended across widespread scalp regions rather than being confined to sensorimotor areas. Whether this broader spatial effect reflects genuine large-scale neural dynamics or methodological bias associated with the analysis procedure remains to be verified by future work. The related limitation also applies to Granger prediction analysis. The same history window was used for both fast and slow movement conditions, as a consequence, the history window contained more trajectory information during faster movements than during slower movements, potentially leading to systematically higher prediction strength in the fast-movement condition. Furthermore, in our design, we cannot distinguish between spectral effects and corresponding (e.g., Granger) results at the key movement frequencies related to visual hand vs target movement signals. Future work should use modified designs, e.g., with variable target frequencies or trajectories, to disentangle these effects, e.g. the Granger prediction at target frequency f0. Finally, although cross-electrode PAC effects based on scalp EEG recordings can be interpreted as reflecting large-scale functional interactions, it should be noted that volume conduction effects and the broad spatial propagation of low-frequency oscillations may lead to signal mixing across electrode sites.

Together, our results identify joint oscillatory activity at key movement frequencies, as well as theta and beta frequencies, as markers of neural processes supporting flexible visuomotor control scaling with the amount of visuo-proprioceptive conflict.

## Acknowledgments

We thank Jan Crusius for providing the eye tracker. Z.W. was supported by the Program of China Scholarship Council (Grant No. 202308310170).

## References

Ablin P, Cardoso J-F, Gramfort A. Faster independent component analysis by preconditioning with Hessian approximations. IEEE Trans Signal Process 66: 4040–4049, 2018.

Ali Bahramisharif, Marcel AJ van Gerven, Erik J Aarnoutse, Manuel R Mercier, Theodore H Schwartz, John J Foxe, Nick F Ramsey, and Ole Jensen. Propagating neocortical gamma bursts are coordinated by traveling alpha waves. Journal of Neuroscience, 33(48):18849–18854, 2013.

Anderson, K. L., & Ding, M. (2011). Attentional modulation of the somatosensory mu rhythm. Neuroscience, 180, 165–180.

Antzoulatos, E. G., & Miller, E. K. (2016). Synchronous beta rhythms of frontoparietal networks support only behaviorally relevant representations. eLife, 5, e17822.

Arnal, L. H., Wyart, V., & Giraud, A. L. (2011). Transitions in neural oscillations reflect prediction errors generated in audiovisual speech. Nature neuroscience, 14(6), 797–801.

Arnal, L. H., Doelling, K. B., & Poeppel, D. (2015). Delta–beta coupled oscillations underlie temporal prediction accuracy. Cerebral Cortex, 25(9), 3077–3085.

Avillac, M., Ben Hamed, S., & Duhamel, J. R. (2007). Multisensory integration in the ventral intraparietal area of the macaque monkey. The Journal of neuroscience, 27(8), 1922–1932.

Balslev, D., Christensen, L. O., Lee, J. H., Law, I., Paulson, O. B., & Miall, R. C. (2004). Enhanced accuracy in novel mirror drawing after repetitive transcranial magnetic stimulation-induced proprioceptive deafferentation. Journal of Neuroscience, 24(43), 9698–9702.

Barnett, L., & Seth, A. K. (2015). Granger causality for state-space models. Physical Review E, 91(4), 040101.

Bastos, A. M., Vezoli, J., Bosman, C. A., Schoffelen, J. M., Oostenveld, R., Dowdall, J. R., De Weerd, P., Kennedy, H., & Fries, P. (2015). Visual areas exert feedforward and feedback influences through distinct frequency channels. Neuron, 85(2), 390–401.

Bauer, M., Kennett, S., Driver, J., 2012. Attentional selection of location and modality in vision and touch modulates low-frequency activity in associated sensory cortices. J. Neurophysiol. 107 (9), 2342–2351. doi: 10.1152/jn.00973.2011

Bauer, M., Oostenveld, R., Peeters, M., Fries, P., 2006. Tactile spatial attention enhances gamma-band activity in somatosensory cortex and reduces low-frequency activity in Parieto-occipital areas. J. Neurosci. 26 (2), 490–501. doi: 10.1523/JNEUROSCI.5228-04.2006

Bauer, M., Stenner, M.-.P., Friston, K., Dolan, R.J., 2014. Attentional modulation of alpha/beta and gamma oscillations reflect functionally distinct processes. J. Neurosci. 34 (48), 16117–16125. doi: 10.1523/JNEUROSCI.3474-13.2014

Bernier, P.-.M., Burle, B., Vidal, F., Hasbroucq, T., Blouin, J., 2009. Direct evidence for cortical suppression of somatosensory afferents during visuomotor adaptation. Cereb. Cortex 19 (9), 2106–2113. doi: 10.1093/cercor/bhn233

Bolt, N. K., & Loehr, J. D. (2024). Motor-related cortical oscillations distinguish one’s own from a partner’s contributions to a joint action. Biological Psychology, 190, 108804.

Botvinick, M., & Cohen, J. (1998). Rubber hands ‘feel’touch that eyes see. Nature, 391(6669), 756–756.

Buschman TJ, Miller EK. Top-down versus bottom-up control of attention in the prefrontal and posterior parietal cortices. Science. 2007;315(5820):1860–2. pmid:17395832

Buschman TJ, Denovellis EL, Diogo C, Bullock D, Miller EK. Synchronous oscillatory neural ensembles for rules in the prefrontal cortex. Neuron 76: 838–846, 2012.

Buzsáki G, Draguhn A. Neuronal Oscillations in Cortical Networks. Science 304: 1926–1929, 2004.

Chacko, R. V., Kim, B., Jung, S. W., Daitch, A. L., Roland, J. L., Metcalf, N. V., … & Leuthardt, E. C. (2018). Distinct phase-amplitude couplings distinguish cognitive processes in human attention. NeuroImage, 175, 111–121.

Clayton, M. S., Yeung, N., & Kadosh, R. C. (2015). The roles of cortical oscillations in sustained attention. Trends in cognitive sciences, 19(4), 188–195.

Cohen, M. X., and T. H. Donner. 2013. “Midfrontal Conflict-Related Theta-Band Power Reflects Neural Oscillations That Predict Behavior.” Journal of Neurophysiology 110, no. 12: 2752–2763.

Cohen, M. X. (2014). Analyzing neural time series data: theory and practice. MIT press.

Coll, M. P., Press, C., Hobson, H., Catmur, C., & Bird, G. (2017). Crossmodal classification of mu rhythm activity during action observation and execution suggests specificity to somatosensory features of actions. Journal of Neuroscience, 37(24), 5936–5947.

Combrisson, E., Nest, T., Brovelli, A., Ince, R. A., Soto, J. L., Guillot, A., & Jerbi, K. (2020). Tensorpac: An open-source Python toolbox for tensor-based phase-amplitude coupling measurement in electrophysiological brain signals. PLoS computational biology, 16(10), e1008302.

De Cheveigné A. ZapLine: a simple and effective method to remove power line artifacts. NeuroImage 207: 116356, 2020.

Delorme A, Makeig S. EEGLAB: an open source toolbox for analysis of single-trial EEG dynamics including independent component analysis. J Neurosci Methods 134: 9–21, 2004.

Drew, A., Gonzalez, J. S. S., Soto-Faraco, S., & Torralba-Cuello, M. (2026). Binocular Rivalry: Evaluating the Role of Theta Power as a Neural Index of Conflict. European Journal of Neuroscience, 63(1), e70372.

Faul, F., Erdfelder, E., Lang, A. G., & Buchner, A. (2007). G* Power 3: A flexible statistical power analysis program for the social, behavioral, and biomedical sciences. Behavior research methods, 39(2), 175–191.

Foxe, J. J., & Snyder, A. C. (2011). The role of alpha-band brain oscillations as a sensory suppression mechanism during selective attention. Frontiers in psychology, 2, 154.

Foxe, J. J., Simpson, G. V., & Ahlfors, S. P. (1998). Parieto-occipital∼ 10Hz activity reflects anticipatory state of visual attention mechanisms. Neuroreport, 9(17), 3929–3933.

Fries P. Rhythms for Cognition: Communication through Coherence. Neuron. 2015 Oct 7;88(1):220–35. doi: 10.1016/j.neuron.2015.09.034. PMID: 26447583; PMCID: PMC4605134.

Fujioka, T., Ross, B., & Trainor, L. J. (2015). Beta-band oscillations represent auditory beat and its metrical hierarchy in perception and imagery. Journal of neuroscience, 35(45), 15187–15198.

Gomez-Ramirez, M., Kelly, S. P., Molholm, S., Sehatpour, P., Schwartz, T. H., & Foxe, J. J. (2011). Oscillatory sensory selection mechanisms during intersensory attention to rhythmic auditory and visual inputs: a human electrocorticographic investigation. Journal of Neuroscience, 31(50), 18556–18567.

González-García, C., Formica, S., Liefooghe, B., & Brass, M. (2020). Attentional prioritization reconfigures novel instructions into action-oriented task sets. Cognition, 194, 104059.

Graziano, M. S. (1999). Where is my arm? The relative role of vision and proprioception in the neuronal representation of limb position. Proceedings of the National Academy of Sciences, 96(18), 10418–10421.

Haegens, S., Händel, B. F., & Jensen, O. (2011). Top-down controlled alpha band activity in somatosensory areas determines behavioral performance in a discrimination task. Journal of neuroscience, 31(14), 5197–5204.

Haegens, S., Luther, L., Jensen, O., 2012. Somatosensory anticipatory alpha activity increases to suppress distracting input. J. Cogn. Neurosci. 24 (3), 677–685. doi: 10.1162/jocn_a_00164

Hari, R. (2006). Action–perception connection and the cortical mu rhythm. Progress in brain research, 159, 253–260.

He, Y., Steines, M., Sammer, G., Nagels, A., Kircher, T., & Straube, B. (2018). Action-related speech modulates beta oscillations during observation of tool-use gestures. Brain Topography, 31(5), 838–847.

Jensen, O., & Mazaheri, A. (2010). Shaping functional architecture by oscillatory alpha activity: gating by inhibition. Frontiers in human neuroscience, 4, 186.

Jones, S. R., Kerr, C. E., Wan, Q., Pritchett, D. L., Hämäläinen, M., & Moore, C. I. (2010). Cued spatial attention drives functionally relevant modulation of the mu rhythm in primary somatosensory cortex. Journal of Neuroscience, 30(41), 13760–13765.

Kaltenmaier, A., Davis, M. H., & Press, C. (2025). Fixed and flexible perceptual rhythms. Trends in Cognitive Sciences.

Keil, J., Pomper, U., & Senkowski, D. (2016). Distinct patterns of local oscillatory activity and functional connectivity underlie intersensory attention and temporal prediction. Cortex, 74, 277–288.

Kelly, E. F., & Folger, S. E. (1999). EEG evidence of stimulus-directed response dynamics in human somatosensory cortex. Brain research, 815(2), 326–336.

Kelso, J. A., Cook, E., Olson, M. E., & Epstein, W. (1975). Allocation of attention and the locus of adaptation tp displaced vision. Journal of Experimental Psychology: Human Perception and Performance, 1(3), 237.

Klug M, Kloosterman NA. Zapline-plus: a zapline extension for automatic and adaptive removal of frequency-specific noise artifacts in M/EEG. Hum Brain Mapp 43: 2743–2758, 2022.

Kothe, C. A., & Makeig, S. (2013). BCILAB: a platform for brain–computer interface development. Journal of neural engineering, 10(5), 056014.

Kulashekhar, S., Pekkola, J., Palva, J. M., & Palva, S. (2016). The role of cortical beta oscillations in time estimation. Human brain mapping, 37(9), 3262–3281.

Kumagai, K. (2021). https://www.sciencedirect.com/science/article/pii/S1063458421006695. Osteoarthritis and cartilage, 29(7), 1020–1028.

Lebar, N., Danna, J., Moré, S., Mouchnino, L., Blouin, J., 2017. On the neural basis of sensory weighting: alpha, beta and gamma modulations during complex movements. Neuroimage 150, 200–212. doi: 10.1016/j.neuroimage.2017.02.043

Limanowski, J., & Friston, K. (2020). Attentional modulation of vision versus proprioception during action. Cerebral Cortex, 30(3), 1637–1648.

Limanowski, J., Litvak, V., & Friston, K. (2020). Cortical beta oscillations reflect the contextual gating of visual action feedback. NeuroImage, 222, 117267.

Limanowski J. Precision control for a flexible body representation. Neurosci Biobehav Rev. 2022 Mar;134:104401. doi: 10.1016/j.neubiorev.2021.10.023. Epub 2021 Nov 1. PMID: 34736884.

Lionel Barnett and Anil K. Seth. Granger causality for state-space models. Physical Review E, 91(4):040101, 2015.

Lloyd, D. M. (2007). Spatial limits on referred touch to an alien limb may reflect boundaries of visuo-tactile peripersonal space surrounding the hand. Brain and cognition, 64(1), 104–109.

Lundqvist, M., Herman, P., Warden, M. R., Brincat, S. L., & Miller, E. K. (2018). Gamma and beta bursts during working memory readout suggest roles in its volitional control. Nature communications, 9(1), 394.

Lundqvist, M., Bastos, A. M., & Miller, E. K. (2020). Preservation and changes in oscillatory dynamics across the cortical hierarchy. Journal of cognitive neuroscience, 32(10), 2024–2035.

Medendorp, W. P., & Heed, T. (2019). State estimation in posterior parietal cortex: Distinct poles of environmental and bodily states. Progress in neurobiology, 183, 101691.

Michalareas, G., Vezoli, J., Van Pelt, S., Schoffelen, J. M., Kennedy, H., & Fries, P. (2016). Alpha-beta and gamma rhythms subserve feedback and feedforward influences among human visual cortical areas. Neuron, 89(2), 384–397.

Miller, E. K., & Buschman, T. J. (2013). Brain rhythms for cognition and consciousness. Neurosciences and the Human Person: New Perspectives on Human Activities, 121.

Miller, K. J., Hermes, D., Honey, C. J., Hebb, A. O., Ramsey, N. F., Knight, R. T., … & Fetz, E. E. (2012). Human motor cortical activity is selectively phase-entrained on underlying rhythms.

Morillon, B., & Baillet, S. (2017). Motor origin of temporal predictions in auditory attention. Proceedings of the National Academy of Sciences, 114(42), E8913–E8921.

Murray, A., Zerroug, Y., Soulières, I., & Saint-Amour, D. (2025). The Role of Fronto-Central Theta Oscillations in Inter-Sensory Selective Attention. Psychophysiology, 62(4), e70055.

Nobre, A. C., Correa, A., & Coull, J. T. (2007). The hazards of time. Current opinion in neurobiology, 17(4), 465–470.

Nordin, A. D., Hairston, W. D., & Ferris, D. P. (2019). Faster gait speeds reduce alpha and beta EEG spectral power from human sensorimotor cortex. IEEE Transactions on Biomedical Engineering, 67(3), 842–853.

Oostenveld R, Fries P, Maris E, Schoffelen J-M. FieldTrip: open source software for advanced analysis of MEG, EEG, and invasive electrophysiological data. Comput Intell Neurosci 2011: 156869, 2011.

Pollok, B., Gross, J., Kamp, D., & Schnitzler, A. (2008). Evidence for anticipatory motor control within a cerebello-diencephalic-parietal network. Journal of cognitive neuroscience, 20(5), 828–840.

Pion-Tonachini, L., Kreutz-Delgado, K., & Makeig, S. (2019). ICLabel: An automated electroencephalographic independent component classifier, dataset, and website. NeuroImage, 198, 181–197

Press, C., Cook, J., Blakemore, S. J., & Kilner, J. (2011). Dynamic modulation of human motor activity when observing actions. Journal of Neuroscience, 31(8), 2792–2800.

Rohe, T., & Noppeney, U. (2018). Reliability-weighted integration of audiovisual signals can be modulated by top-down attention. eneuro, 5(1).

Saleh, M., Reimer, J., Penn, R., Ojakangas, C. L., & Hatsopoulos, N. G. (2010). Fast and slow oscillations in human primary motor cortex predict oncoming behaviorally relevant cues. Neuron, 65(4), 461–471.

Senkowski, D., & Engel, A. K. (2024). Multi-timescale neural dynamics for multisensory integration. Nature Reviews Neuroscience, 25(9), 625–642.

Senkowski, D., Schneider, T. R., Foxe, J. J., & Engel, A. K. (2008). Crossmodal binding through neural coherence: implications for multisensory processing. Trends in neurosciences, 31(8), 401–409

Sober, S. J., & Sabes, P. N. (2005). Flexible strategies for sensory integration during motor planning. Nature neuroscience, 8(4), 490–497.

Spitzer, B., Haegens, S., 2017. Beyond the status quo: a role for beta oscillations in endogenous content (re)activation. ENeuro 4 (4), 2017. doi: 10.1523/ENEURO.0170-17.2017, ENEURO.0170-17

Spooner, R. K., Wiesman, A. I., Proskovec, A. L., Heinrichs - Graham, E., & Wilson, T. W. (2020). Prefrontal theta modulates sensorimotor gamma networks during the reorienting of attention. Human Brain Mapping, 41(2), 520–529.

Thomas S. Binns, Adam Li, Eric Larson, Daniel McCloy, Alex Rockhill, Qianliang Li, Santeri Ruuskanen, Alexander Kroner, Alexandre Gramfort, Giovanni Franco Gabriel Marraffini, Jonathan Shor, Kenji Marshall, Mohammad Orabe, Moritz Gerster, Richard M. Köhler, Sam Steingold, and Sezan Mert. MNE-Connectivity. 2026.

Thut, G., Nietzel, A., Brandt, S. A., & Pascual-Leone, A. (2006). α-Band electroencephalographic activity over occipital cortex indexes visuospatial attention bias and predicts visual target detection. Journal of neuroscience, 26(37), 9494–9502.

Toma, K., Mima, T., Matsuoka, T., Gerloff, C., Ohnishi, T., Koshy, B., … & Hallett, M. (2002). Movement rate effect on activation and functional coupling of motor cortical areas. Journal of neurophysiology, 88(6), 3377–3385.

Trajkovic, J., Veniero, D., Hanslmayr, S., Palva, S., Cruz, G., Romei, V., & Thut, G. (2025). Top-down and bottom-up interactions rely on nested brain oscillations to shape rhythmic visual attention sampling. PLoS Biology, 23(4), e3002688.

Ullsperger, M., Danielmeier, C., & Jocham, G. (2014). Neurophysiology of performance monitoring and adaptive behavior. Physiological reviews, 94(1), 35–79.

Van Beers, R.J., Sittig, A.C. & van der Gon, J.J.D. Integration of proprioceptive and visual position-information: an experimentally supported model. J. Neurophysiol. 81, 1355–1364 (1999)

Van Ede, F., Jensen, O., & Maris, E. (2010). Tactile expectation modulates pre-stimulus β-band oscillations in human sensorimotor cortex. Neuroimage, 51(2), 867–876.

Van Ede, F., De Lange, F., Jensen, O., & Maris, E. (2011). Orienting attention to an upcoming tactile event involves a spatially and temporally specific modulation of sensorimotor alpha-and beta-band oscillations. Journal of Neuroscience, 31(6), 2016–2024.

Van Moorselaar, D., & Slagter, H. A. (2020). Inhibition in selective attention. Annals of the New York Academy of Sciences, 1464(1), 204–221.

Vigh, G., & Limanowski, J. (2025). Baseline dependent differences in the perception of changes in visuomotor delay. Frontiers in Human Neuroscience, 18, 1495592.

Wang, P., & Limanowski, J. (2023). Phasic modulation of beta power at movement-related frequencies during visuomotor conflict. Journal of Neurophysiology, 130(5), 1367–1372.

Watanabe, T., Mima, T., Shibata, S., & Kirimoto, H. (2021). Midfrontal theta as moderator between beta oscillations and precision control. Neuroimage, 235, 118022.

Weisz, N., Hartmann, T., Müller, N., Lorenz, I., & Obleser, J. (2011). Alpha rhythms in audition: cognitive and clinical perspectives. Frontiers in psychology, 2, 73.

Wendiggensen, P., Prochnow, A., Pscherer, C., Münchau, A., Frings, C., & Beste, C. (2023). Interplay between alpha and theta band activity enables management of perception-action representations for goal-directed behavior. Communications Biology, 6(1), 494.

Wolpert, D. M., Goodbody, S. J., & Husain, M. (1998). Maintaining internal representations: the role of the human superior parietal lobe. Nature neuroscience, 1(6), 529–533.

Zavala, B., A. Jang, M. Trotta, C. I. Lungu, P. Brown, and K. A. Zaghloul. 2018. “Cognitive Control Involves Theta Power Within Trials and Beta Power Across Trials in the Prefrontal-Subthalamic Network.” Brain 141, no. 12: 3361–3376.

Zeller, D., Litvak, V., Friston, K. J., & Classen, J. (2015). Sensory processing and the rubber hand illusion—an evoked potentials study. Journal of Cognitive Neuroscience, 27(3), 573–582.

