## Supplementary material for "Oscillatory correlates of visuomotor control under varying amount of feedback delay"

#### *Grasp range*

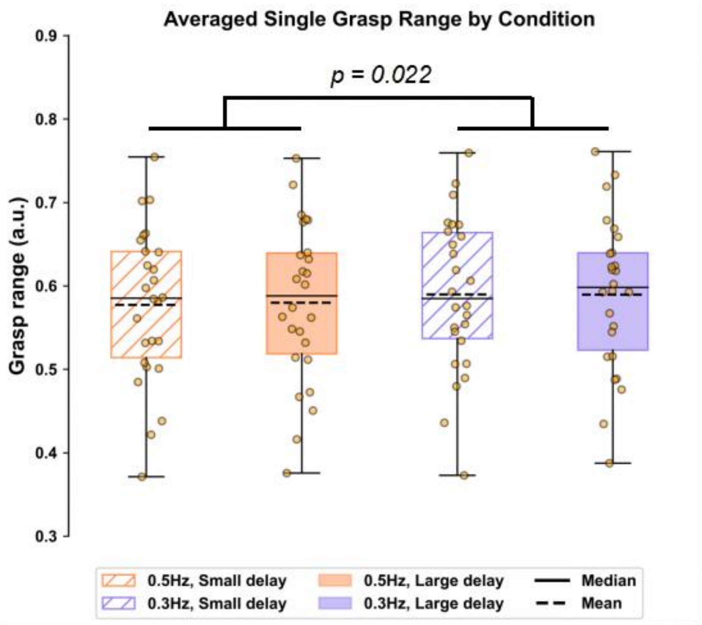


**Figure S1.** Movement amplitude/grasp range (maximum hand opening to minimum hand closing) across experimental conditions. Linear mixed-effects model (*Grasp range ~ Target Frequency * Delay + (1|Participant) )* revealed a significant main effect of target frequency (*F(1,75)*=5.43, *p*=0.022, $dz$=-0.36). Participants shown a slightly smaller grasp range at 0.5 Hz (mean ± SEM: 0.579 ± 0.018) than target frequency 0.3 Hz condition (0.590 ± 0.018). Although this difference was statistically significant, it only amounted to about 1.9% of the overall movement amplitude, rendering it practically negligible. Most importantly, delay had no significant effect on movement amplitudes (*F(1,75)*=0.069, *p*=0.79, $dz$=0.047).

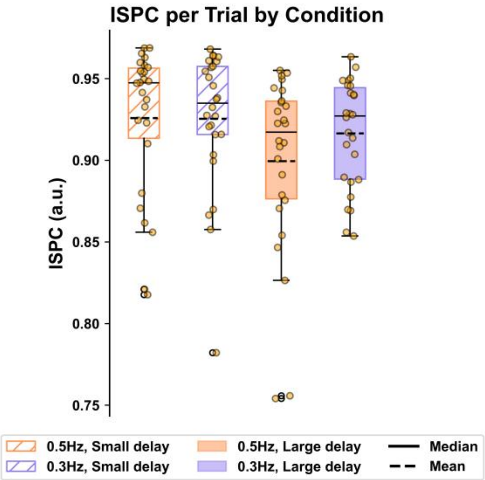


**Figure S2.** Average inter-site phase clustering (ISPC) for each experimental condition. ISPC measures the consistency of the phase relationship between two trajectories across time regardless of the phase shift error. The single trial ISPC was estimated through $ISPC = |n^{-1}\sum_{t=1}^{n} e^{i\Delta\emptyset_t}|$, where n is the number of time samples and $\Delta\emptyset_t$ is the phase difference between the VH and the target. The phase time courses of the VH and target trajectories were obtained from Hilbert transformation. For each participant, each condition, the ISPC was averaged across 15 trials. Higher average ISPC value indicates more continuous and stable tracking, with a value of 1 reflecting perfect phase locking. Overall, participants tracked the target smoothly across all conditions (ISPC group means > 0.9). Tracking performed more stable at small > large delays ($F(1,75)$=14.43, $p$=0.00029; $dz$=−0.67).

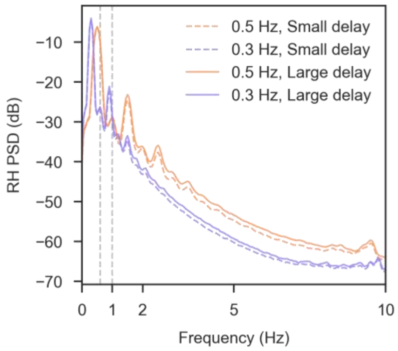


**Figure S3.** Power spectral density (PSD) of hand movement trajectory averaged across participants. Dash lines marked the f1- peak frequency at 0.6Hz and 1 Hz. The dominant f0- peaks in spectrum were in alignment with the target frequency at 0.3 Hz and 0.5Hz for the corresponded experimental condition. Smaller first harmonic components (f1) were also present in the spectrum.

### *PSD topology*

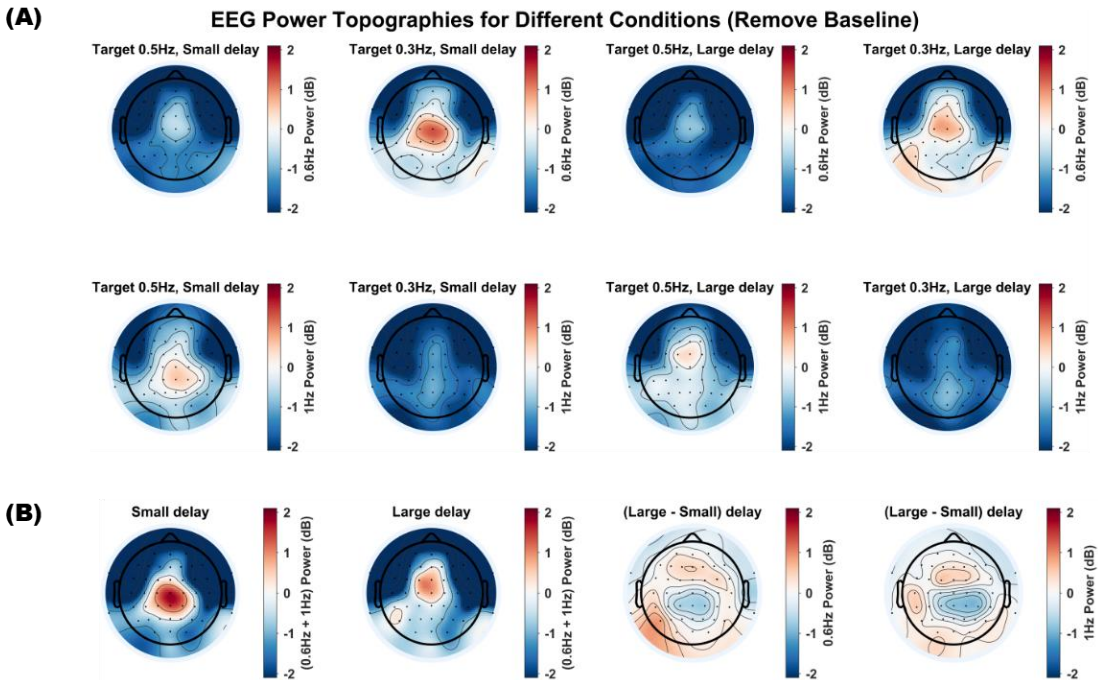


**Figure S4.** (A) Topographical distribution of EEG power at 0.6 Hz (first row) and 1 Hz (second row ) across experimental conditions, baseline corrected (relative to a pre-movement fixation only period, see Methods). (B) Visualization of the delay effect topology on first harmonic (0.6 Hz and 1 Hz) power.

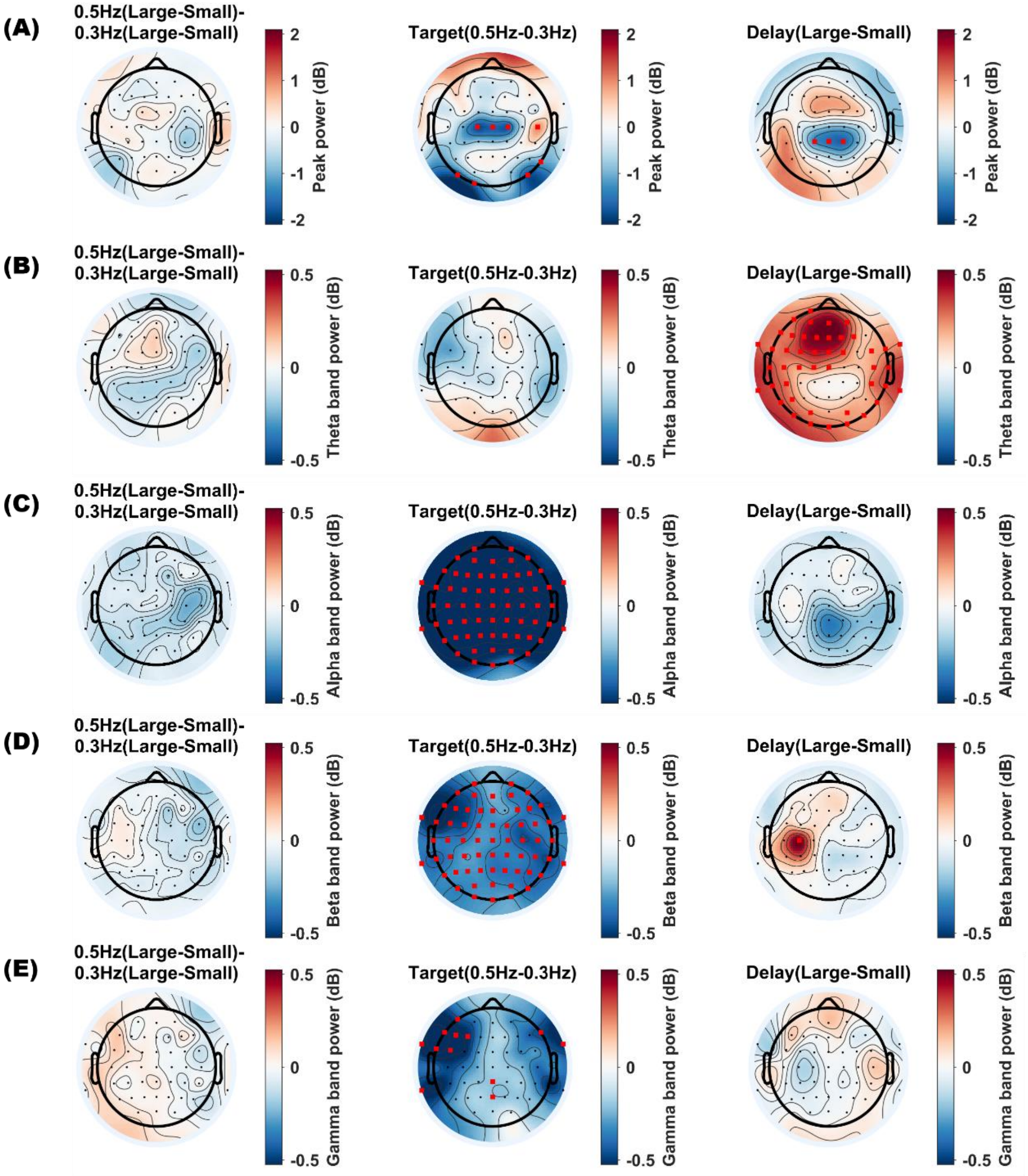


**Figure S5.** Linear mixed-effects model (*Power ~ Target Frequency * Delay + (1|Participant)*) results for EEG sensor-level power across frequency bands. (A) Peak power at the first harmonic of the target frequency, already shown in main text. (B) Theta (4–7 Hz), (C) alpha (8–13 Hz), (D) beta (14–30 Hz) bands, and (E) gamma (31-70 Hz). Significant channels were marked in red. In addition to delay, we observed significant effects of target frequency; however, a potential limitation is the use of different epoch lengths for the 0.5 Hz and 0.3 Hz conditions in the continuous wavelet transform. Differences in epoch duration can influence power estimates, such that longer epochs (target 0.3 Hz) may yield higher alpha (8–12 Hz) and beta (13–30 Hz) band power. Therefore, the stronger power observed in longer epochs may have resulted from analytical confounds instead of reflecting meaningful neurophysiological differences. Please note, however, that our primary focus was on the orthogonal delay effect—the key factor underlying visuo-proprioceptive conflict in our design.

### *Granger prediction*

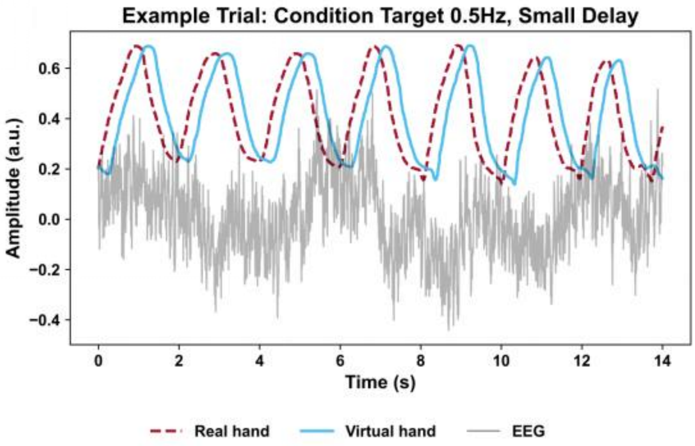


**Figure S6.** Example single trial (0.5 Hz, small delay) illustrating the concatenation of seven grasping movements with simultaneous EEG recordings at CPz. These data were used to estimate Granger-prediction relationships among the real hand movement (red dashed line), the virtual hand movement (blue solid line), and EEG activity (gray line).

**Permutation-based sanity check for the directional Granger prediction**

To verify that the observed Granger-prediction effects reflected genuine directional coupling rather than random fluctuations, we performed a permutation test at the single-subject level. Specifically, we used non-overlapping trial segments (15 trials, each containing seven grasping cycles) to compute the observed mean Granger-prediction value. Trial labels were then randomly permuted 30 times to generate a null distribution of Granger-prediction means. From this null distribution, we computed the mean and standard deviation and estimated a one-tailed p-value at the first harmonic frequency using a Gaussian approximation, with the lower bound capped at 1/31 ($\approx$ 0.03) to reflect the finite number of permutations. At the group level, individual p-values were combined using Fisher's method. The combined statistics were evaluated separately for the 1 Hz and 0.6 Hz directions (one-tailed tests). Across all conditions, the Granger prediction from EEG to hand at the first harmonic frequency (1 Hz or 0.6 Hz correspondingly) was significant (Table S1). This analysis served as a sanity check, confirming consistent directional Granger-prediction effects across participants.

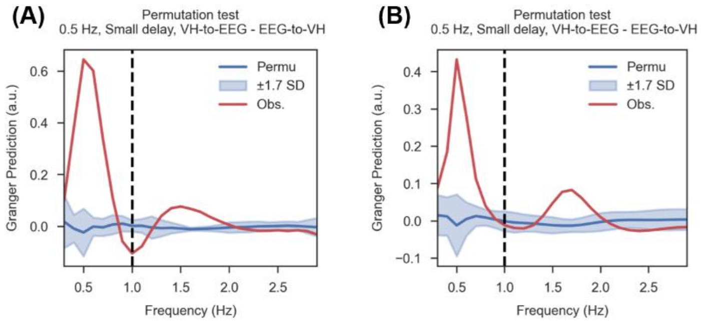


**Figure S7.** Example of the observed directional Granger prediction (Obs., red line) and the permuted directional Granger prediction (Permutation mean, blue line) for a single subject in the target 0.5 Hz / small delay condition. The blue shading indicates the one-tailed significance threshold (1.7 standard deviations (SD), corresponding to $p < 0.05$). The dashed line marks the first harmonic response of the 0.5 Hz target at 1 Hz. (A) history window N=20 samples; (B) history window N=5 samples.

**Table S1.** Group-level Fisher's combined significance across conditions (history window N=20 samples). Individual one-tailed p-values (from subject-level Granger-prediction analyses) were combined using Fisher's method. Reported are the resulting $X^2$ statistics (df = 52) and corresponding uncorrected p-values for EEG to real hand (EEG → RH) and EEG to visual hand (EEG → VH) predictions under each combination of target frequency and delay.

| Target | Delay | Prediction direction | $X^2(52)$ | p value (uncor.) |
|---|---|---|---|---|
| 0.5 Hz | Small delay | EEG->RH | 132.30 | p = 6.23e-9 |
| | | EEG->VH | 155.09 | p = 3.44e-12 |
| 0.3 Hz | Small delay | EEG->RH | 133.91 | p = 3.74e-9 |
| | | EEG->VH | 163.94 | p = 1.61e-13 |
| 0.5 Hz | Large delay | EEG->RH | 127.65 | p = 2.66e-8 |
| | | EEG->VH | 165.28 | p = 1.01e-13 |
| 0.3 Hz | Large delay | EEG->RH | 155.44 | p = 3.05e-12 |
| | | EEG->VH | 174.49 | p = 3.82e-15 |

**Table S2.** Group-level Fisher's combined significance across conditions (history window N=5 samples). Individual one-tailed p-values (from subject-level Granger-prediction analyses) were combined using Fisher's method. Reported are the resulting $X^2$ statistics (df = 52) and corresponding uncorrected p-values for EEG to real hand (EEG → RH) and EEG to visual hand (EEG → VH) predictions under each combination of target frequency and delay.

| Target | Delay | Prediction direction | $X^2(52)$ | p value (uncor.) |
|---|---|---|---|---|
| 0.5 Hz | Small delay | EEG->RH | 114.33 | p = 1.42e-6 |
| | | EEG->VH | 128.99 | p = 1.75e-8 |
| 0.3 Hz | Small delay | EEG->RH | 52.05 | p = 0.47 |
| | | EEG->VH | 67.99 | p = 0.068 |
| 0.5 Hz | Large delay | EEG->RH | 94.18 | p = 0.00031 |
| | | EEG->VH | 136.98 | p = 1.41e-9 |
| 0.3 Hz | Large delay | EEG->RH | 70.58 | p = 0.044 |
| | | EEG->VH | 80.91 | p = 0.0063 |

**Table S3.** One-sample t-tests of the directional spectral Granger prediction (EEG → hand) at the first harmonic frequency (f1) across experimental conditions.

| Conditions | EEG-> | Mean (SD) | t(25) | FDR corrected p |
|---|---|---|---|---|
| 0.5Hz, small delay | VH | -0.117 (0.058) | -10.33 | 2.23e-10 |
| 0.5Hz, large delay | VH | -0.118 (0.048) | -12.68 | 5.84e-12 |
| 0.3Hz, small delay | VH | -0.084 (0.030) | -14.13 | 1.61e-12 |
| 0.3Hz, large delay | VH | -0.082 (0.032) | -13.10 | 4.26e-12 |
| 0.5Hz, small delay | RH | -0.086 (0.056) | -7.92 | 3.04e-8 |
| 0.5Hz, large delay | RH | -0.069 (0.045) | -7.89 | 3.04e-8 |
| 0.3Hz, small delay | RH | -0.060 (0.028) | -11.14 | 6.92e-11 |
| 0.3Hz, large delay | RH | -0.053 (0.026 ) | -10.35 | 2.23e-10 |

**Table S4.** One-sample t-tests of the directional spectral Granger prediction (EEG → hand) at the first harmonic frequency (f0) across experimental conditions.

| Conditions | EEG-> | Mean (SD) | t(25) | FDR corrected p |
|---|---|---|---|---|
| 0.5Hz, small delay | VH | 0.742 (0.121) | 31.23 | 3.01e-21 |
| 0.5Hz, large delay | VH | 0.729 (0.093) | 40.04 | 9.03e-24 |
| 0.3Hz, small delay | VH | 0.302 (0.108) | 14.30 | 1.83e-13 |
| 0.3Hz, large delay | VH | 0.271 (0.094) | 14.63 | 1.48e-13 |
| 0.5Hz, small delay | RH | 0.735 (0.093) | 40.23 | 9.03e-24 |
| 0.5Hz, large delay | RH | 0.722 (0.086) | 42.79 | 5.27e-24 |
| 0.3Hz, small delay | RH | 0.291 (0.129) | 11.49 | 1.80e-11 |
| 0.3Hz, large delay | RH | 0.274 (0.096) | 14.50 | 1.51e-13 |

**Table S5.** Three-way repeated measures ANOVA testing how experimental conditions modulate EEG (CP) prediction direction dominance of real and virtual hand at the first harmonic frequency (f1). Factors: Hand (RH, VH) × Target Frequency (0.5 Hz, 0.3 Hz) × Delay (small, large).

| | F(1,25) | uncorr. p |
|---|---|---|
| **Target frequency** | **16.64** | **0.00040** |
| Delay | 1.19 | 0.29 |

| | | |
|---|---|---|
| **Hand** | **84.37** | **1.73e-9** |
| Target frequency : Delay | 0.079 | 0.78 |
| **Target frequency : Hand** | **6.18** | **0.020** |
| **Delay : Hand** | **6.45** | **0.018** |
| Target frequency : Delay : Hand | 1.92 | 0.18 |

**Table S6.** Three-way repeated measures ANOVA testing how experimental conditions modulate EEG (CP) prediction direction dominance of real and virtual hand at the target frequency (f0). Factors: Hand (RH, VH) × Target Frequency (0.5 Hz, 0.3 Hz) × Delay (small, large).

| | F(1,25) | uncorr. p |
|---|---|---|
| **Target frequency** | **1288.34** | **4.98e-23** |
| **Delay** | **6.21** | **0.0197** |
| Hand | 0.46 | 0.51 |
| Target frequency : Delay | 0.32 | 0.58 |
| Target frequency : Hand | 0.027 | 0.87 |
| Delay : Hand | 0.35 | 0.56 |
| Target frequency : Delay : Hand | 0.37 | 0.55 |

### *Gaze fixation*

**Table S7.** Results of the linear mixed-effects model testing whether gaze fixation differed across experimental conditions. A linear mixed-effects model (n=18), similar as in Sections 2.4.1, 2.4.2 & 2.4.4 was conducted to assess whether the mean eye-gaze distance from the central fixation point varied with target frequency, delay, or their interaction. No significant effects were found.

| | F | df1 | df2 | p-value | dz |
|---|---|---|---|---|---|
| Target frequency | 0.00152 | 1 | 51 | 0.990 | 0.0030 |
| Delay | 0.805 | 1 | 51 | 0.374 | 0.22 |
| Target frequency: Delay | 0.00265 | 1 | 51 | 0.959 | -0.011 |
